# Demystifying “drop-outs” in single cell UMI data

**DOI:** 10.1101/2020.03.31.018911

**Authors:** Tae Kim, Xiang Zhou, Mengjie Chen

**Affiliations:** Department of Statistics, University of Chicago; Department of Biostatistics, University of Michigan; Department of Human Genetics and Department of Medicine, University of Chicago

## Abstract

Analysis of scRNA-seq data has been challenging particularly because of excessive zeros observed in UMI counts. Prevalent opinions are that many of the detected zeros are “drop-outs” that occur during experiments and that those zeros should be accounted for through procedures such as normalization, variance stabilization, and imputation. Here, we extensively analyze publicly available UMI datasets and challenge the existing scRNA-seq workflows. Our results strongly suggest that resolving cell-type heterogeneity should be the foremost step of the scRNA-seq analysis pipeline because once cell-type heterogeneity is resolved, “drop-outs” disappear. Additionally, we show that the simplest parametric count model, Poisson, is sufficient to fully leverage the biological information contained in the UMI data, thus offering a more optimistic view of the data analysis. However, if the cell-type heterogeneity is not appropriately taken into account, pre-processing such as normalization or imputation becomes inappropriate and can introduce unwanted noise. Inspired by these analyses, we propose a zero inflation test that can select gene features contributing to cell-type heterogeneity. We integrate feature selection and clustering into iterative pre-processing in our novel, efficient, and straightforward framework for UMI analysis, HIPPO (Heterogeneity Inspired Pre-Processing tOol). HIPPO leads to downstream analysis with much better interpretability than alternatives in our comparative studies.

## Main

Droplet-based single cell RNA-sequencing (scRNA-seq) methods have changed the landscape of genomics research in complex biological systems [19, 21, 36, 37] by producing single cell resolution data at affordable costs. In the state-of-the-arts protocols, a step called barcoding unique molecular identifiers (UMI) has been introduced to remove amplification bias and further improve data quality [16]. Some literature [5, 29] suggests that barcoding leads to a different data structure from read count data structure but many tools remain to not acknowledge the difference between the count data produced with and without barcoding.

Many pipelines have been built for scRNA-seq UMI data analysis. Despite subtle differences in these pipelines, the general order of a scRNA-seq analysis is as follows: quality control (filtering), cleaning (normalization, imputation, de-noising, batch-correction, etc.), feature selection which often involves dimension reduction, and downstream analysis such as clustering and lineage analysis. In this paper, we do not discuss filtering and focus on the later three steps. First, the challenge of scRNA-seq data cleaning has led to the development of a wide array of tools. Some methods adjust for sequencing depths using size factors [4, 32]. Some impute the reads directly using a zero inflated model, to reduce the noise from drop-outs [10]. Some try to de-noise the entire data set by fitting parametric models, where one example is sctransform that uses the residuals from negative binomial regression [11] and another example is SAVER that uses Poisson LASSO regression [13]. Despite the diversity of proposed methods, the general consensus has been reached to use one of the following distributions to model the counts: Poisson, Negative Binomial, or Zero inflated Negative Binomial distribution. Secondly, methods for feature selection have been less controversial. Most tools use some form of gene variance to mean ratio to identify genes that are highly dispersed, where the dispersion level is interpreted as a signal of biological heterogeneity [4, 11, 5]. Another less recognized approach is to use the zeros in the read or UMI counts; genes with inflated zeros are interpreted as biologically important signals [1]. Lastly, after data cleaning and feature selection, the pre-processed data will then be piped into downstream analysis tools for clustering analysis [34, 8, 9], trajectory inference [24, 30], or differential expression analysis [23, 20]. Currently, pre-processing and downstream analysis have been mostly considered as separate and consecutive steps [8, 4, 13, 35].

Here, we present extensive analyses of publicly available UMI data sets that challenge most existing pre-processing tools’ assumption, mainly that pre-processing is a necessary step before feature selection and downstream analysis. Our results suggest that clustering, or resolving the cell heterogeneity, should be the foremost step of the scRNA-seq analysis pipeline, not as part of the downstream analysis. Normalizing or imputing the data set before resolving the heterogeneity can lead to adversary consequences in downstream analysis. Adding to the arguments that the UMI data is much cleaner than the read count data [5, 29], our analyses demonstrate that the simple Poisson distribution is sufficient to fully leverage the biological information contained in the UMI data if the cell-type heterogeneity has been appropriately accounted for. As a result, we provide a new perspective on scRNA-seq data analysis by integrating the pre-processing step and clustering, which was classified as part of the down-stream analysis. The proposed procedures have been implemented in software HIPPO (https://github.com/tk382/HIPPO).

## Results

### Demystifying Drop-outs

We started by exploring zero detection rates in three UMI datasets generated by 10X protocols for both homogeneous and heterogeneous cell populations. Taking a subset of data in Zheng 2017 [36] as created in Freytag 2018 [9] as an example, we computed zero proportions, defined as the proportion of cells with zero counts per gene, across 15,568 genes, in CD19+ B cells, CD4+/CD25 Regulatory T cells, and combined. The obtained statistics were plotted against gene-level average count and were compared with expected zero proportions under the Poisson, Negative Binomial and Zero-Inflated Negative Binomial distribution, respectively (Figure 1 A). For a homogeneous cell population, we observe most genes align well with the expected curve under the Poisson assumption. Few genes can benefit from using the Negative Binomial model to account for extra dispersion from the Poisson, but our results strongly suggest that to model the drop-outs by introducing an extra zero-inflation component by the Zero-Inflated Negative Binomial distribution is unnecessary. For example, in Zheng dataset, 257 genes out of 5,568 genes would benefit from Negative Binomial modeling (p-values pass Bonferroni criterion in likelihood ratio test at 0.05 type I error level), but no gene would benefit from extra zero inflation parameter. The p-values are not calibrated to the uniform distribution because there are many genes that have UMI count of 1 in one cell and 0 in everywhere else, in which case p-value is close to 1 (Figure 1 B). This result shows that drop-outs are within the range of natural Poisson sampling noise in UMI data for a homogeneous cell population, and they do not introduce excessive zero inflation, which is contradictory to prevalent opinions [8, 22, 31, 32]. Extra zero inflation can be measured through a test statistic of z-score that follows standard normal distribution within a completely homogeneous cell population (Methods). Through the following analysis, we find zero proportions are better indicators for cell-type heterogeneity than widely used alternatives, gene variance, coefficient of variation (CV), or dispersion parameter in negative binomial distribution [1] (Figure 1 A).

**Figure 1:**
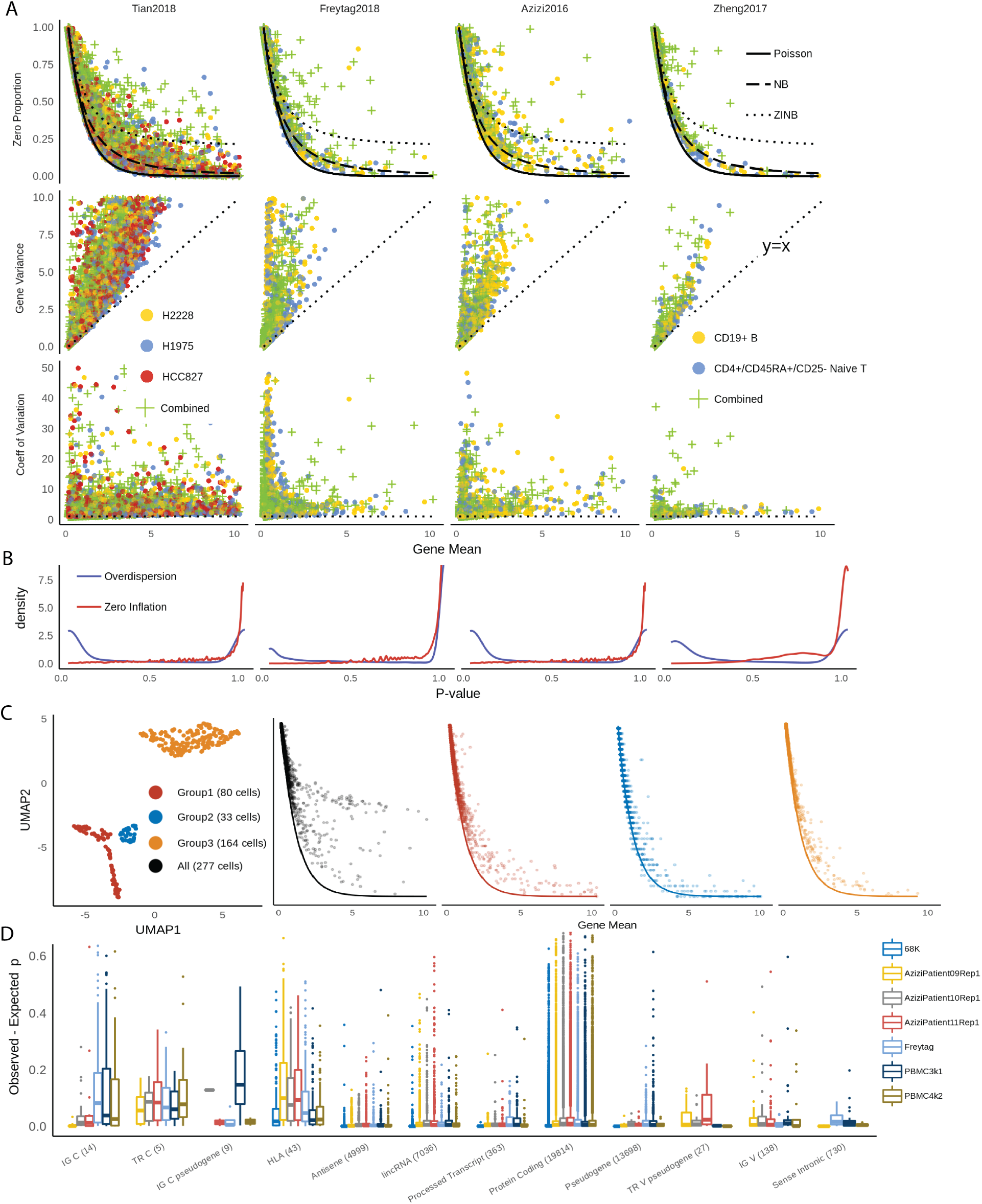
A. Comparisons of zero proportion, gene variance and CV as indicators for cellular heterogeneity in different UMI data sets. B. Distributions of *p*-values from likelihood ratio test for over-dispersion and zero-inflation. C. t-SNE plots of CD34+ cells in Zheng data, and relationship between zero proportions and gene means before (black) and after (colors) clustering of CD34+ cells. D. Distributions of zero inflation in different PBMC data sets. The x-axis labels represent gene types from GENCODE annotations and the number of genes within each type.

Analysis in multiple UMI data sets shows that zero proportions in most genes can be effectively modeled by the Poisson distribution, as more than 95% of absolute z-values are below 2. For mixed cell types, zero proportions considerably deviate from expected values under the Poisson model, as only less than 30% of the genes have z-values below 2. This shows that the zero inflation test is an effective way to find genes that contribute to cellular heterogeneity. On the contrary, gene variance of mixed cell types does not always surpass those of a single cell type. In Zheng data, 62% of the genes had higher variance in pure Naive Cytotoxic cells than in mixed PBMC cells. On average, gene variance is similarly distributed for homogeneous and heterogeneous cell populations (Supplementary Table 2, Supplementary Figure 4). Therefore, the gene variance is rather more of a gene-specific characteristic while being less informative about the characteristics of the entire cell population. CV, on the other hand, suffers from an inherent numerical instability issue when gene mean is close to 0, because when mean is close to 0, CV estimates have high variability. Another popular option is to conduct model selection to assign genes to one of three candidate distributions of Poisson, NB, and ZINB, but measuring over-dispersion also suffers from a similar problem in selecting biologically meaningful genes [6, 5]. When we used statistics from likelihood ratio test and select top genes from the resulting statistics, the selected genes were very different from those selected when we used zero proportion (Figure 2 F). For three data sets of Azizi2016, Zheng2017, and Freytag2018 (median sequencing depth of 4371, 1298, and 2393.5 respectively), the likelihood ratio test selects genes that are overly focused on those with mean close to 0. Intuitively, the dispersion parameter scales with the ratio of gene mean to the gene variance^2^, and in nature very similar to CV. These genes with very low mean are likely to have little information about the cells. Tian2018 data has particularly high sequencing depth (median 107648), so the feature selection behaves differently, but still, the final feature set from two methods differ from one another.

**Table 1:** Zero inflation test statistics of PPBP gene in CD34+ cells in Zheng data before and after clustering into subtypes.

| Cell Population | gene mean | expected $p$ | observed $p$ | $z$-score |
| --- | --- | --- | --- | --- |
| CD34+ | 25.89 | 5.69e-12 | 0.26 | 1838203 |
| Subtype 1 | 0.5625 | 0.57 | 0.91 | 6.19 |
| Subtype 2 | 22.36 | 1.93e-10 | 0 | 0 |
| Subtype 3 | 38.96 | 1.25e-17 | 0 | 0 |

**Table 2:** List of data sets used in the main text and supplementary materials.

| ID | Data Set | Species | Protocol | Year | # Genes | # Cells |
| --- | --- | --- | --- | --- | --- | --- |
| 10x | 5KNeuron | Mouse | 10xGenomics | 2019 | 31053 | 6997 |
| 10x | 10KHeart | Mouse | 10xGenomics | 2018 | 31053 | 7713 |
| GSE111108 [28] | Tian2018 | Human | 10xGenomics | 2018 | 58302 | 925 |
| GSE115189[9] | Freytag2018 | Human | 10xGenomics | 2018 | 58302 | 2590 |
| 10x | 1KNeuron | Mouse | 10xGenomics | 2017 | 31053 | 1206 |
| SRP073767[36] | Zhengmix4eq | Human | 10xGenomics | 2017 | 15568 | 3994 |
| SRP073767 | Zhengmix4uneq | Human | 10xGenomics | 2017 | 16443 | 6498 |
| SRP073767 | Zhengmix8efq | Human | 10xGenomics | 2017 | 15176 | 3994 |
| SRP073767 | PBMC3k | Human | 10xGenomics | 2017 | 58302 | 3250 |
| SRP073767 | PBMC4k | Human | 10xGenomics | 2017 | 58302 | 4292 |
| SRP073767 | PBMC68k | Human | 10xGenomics | 2017 | 32738 | 68579 |
| GSE84133[3] | Baron | Human | inDrop | 2016 | 19097 | 350 |
| GSE114724[2] | AziziPatient09Rep1 | Human | 10xGenomics | 2016 | 33694 | 5287 |
| GSE114724 | AziziPatient09Rep2 | Human | 10xGenomics | 2016 | 33694 | 5256 |
| GSE114724 | AziziPatient10Rep1 | Human | 10xGenomics | 2016 | 33694 | 3349 |
| GSE114724 | AziziPatient11Rep1 | Human | 10xGenomics | 2016 | 33264 | 5287 |
| GSE114724 | AziziPatient11Rep2 | Human | 10xGenomics | 2016 | 33694 | 3696 |
| GSE63473[21] | Macosko | Mouse | Drop-seq | 2015 | 23288 | 44808 |

**Figure 2:**
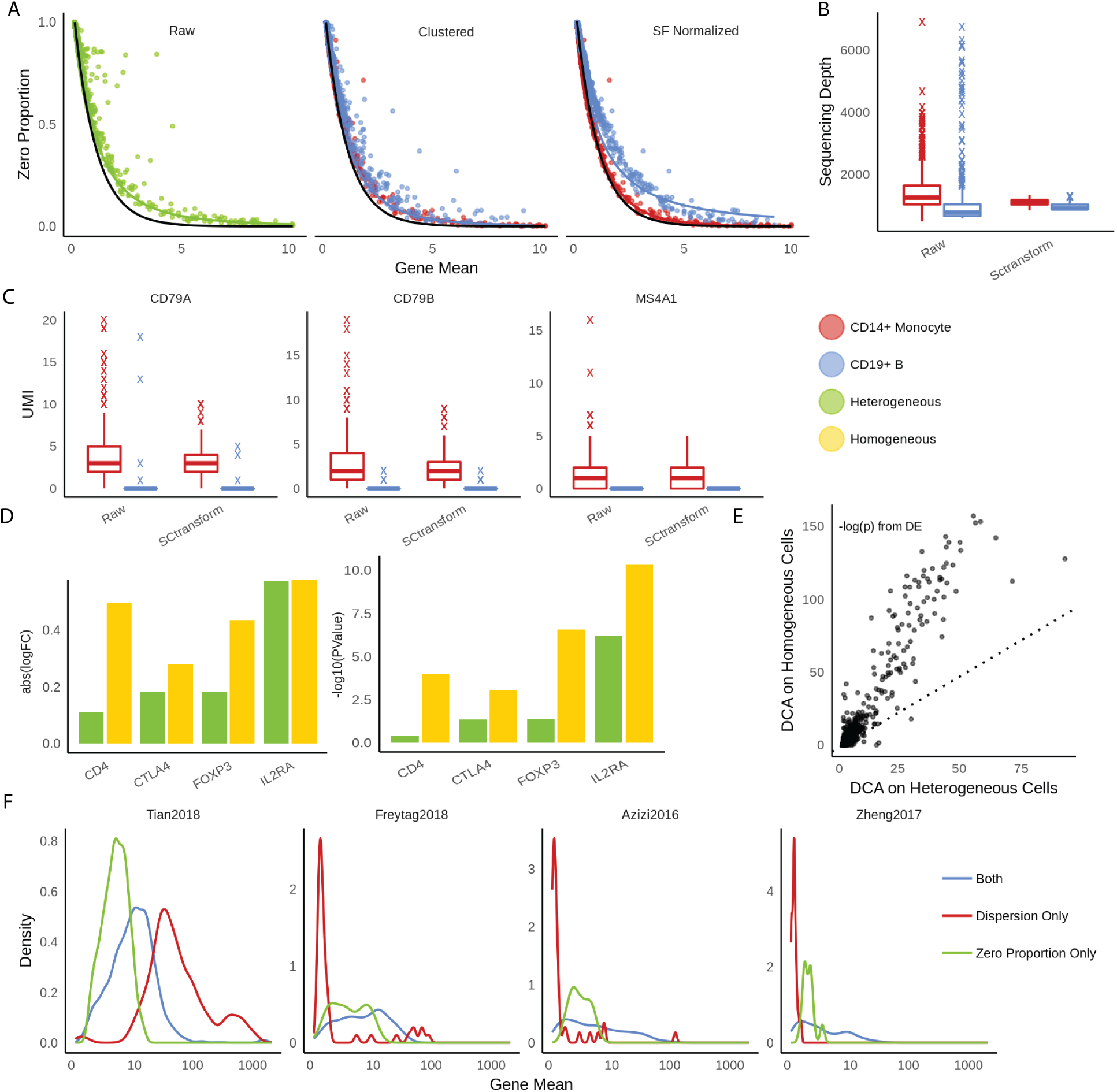
A. Scatterplots between gene means and zero proportions across genes calculated from raw UMI data, clustered data, and data after sequencing-depth normalization, respectively. Fitted line is negative binomial curve. B-C. Evaluations of pre-processing in Sctransform. B. Distributions of sequencing depths across cells in raw UMI data vs. data cleaned by SCtransform. C. Comparisons of three Monocyte markers in raw UMI data vs. data cleaned by SCtransform. D-E. Evaluations of pre-processing in DCA. D. Log fold changes and log p-values from differential expression analysis using the same data set but imputed by DCA as heterogeneous and homogeneous cell populations, respectively. E. The p-values comparisons between two different imputation strategies show the general deflation of biological signals of DCA when applied to heterogeneous cells. F. Comparisons of selected features using likelihood ratio test and zero proportions. Dispersion tends to select the genes that have mean UMI count close to 0.

We expand the data to study all 68,579 cells from Zheng dataset [36]. When we aligned the zero proportions with the expected Poisson curve according to the provided cell type labels: CD14+ Monocyte, CD19+ B, CD34+, CD4+ T Helper2, CD4+/CD25 T Reg, CD45RA+/CD25-Naive T, CD4+/CD45RO+ Memory, CD56+ NK, CD8+ Cytotoxic T, CD8+/CD45RA+ Naive Cytotoxic, and Dendritic cells. Most of these cell types look relatively homogeneous. However, one cell type, CD34+, was particularly noisy with very high zero proportions, indicating cellular heterogeneity (Figure 1 C). Based on the diagnosis from t-SNE plots, we identified three subtypes within the CD34+ cells. The alignment of zero proportions against the Poisson curve was immediately improved according to the inferred subtype labels. This indicates the effectiveness of zero proportions as metrics to evaluate cellular heterogeneity and their potentials to discern cell types.

We further checked how zero proportions could be dispersed from the Poisson distribution for genes with various functional annotations across all PBMC datasets (Figure 1 D). Specifically, we calculated the difference between observed zero proportions and expected proportions (under Poisson) for each functional group using reference data from GENCODE of GRCh38 [12]. The vast majority of genes are categorized as “protein-coding genes”. Their zero proportions cover a wide range from 0 to 0.7, but centered at 0, indicating variability in zero proportions but no systematic inflation of zero proportions. In contrast, immune-related genes are consistently zero-inflated, with the interquartile range as high as 10% to 20%. The enrichment analysis (Supplementary Table 4) shows that immune-related genes have significantly higher proportion of zero-inflated genes compared to genes that are not related to immune function. The top-ranked annotations for zero inflation include IG C genes, TR C genes, and HLA genes. IG C genes are immunoglobulin genes of the constant (C) region, while TR C genes are T cell receptor genes of the constant region, and hence both gene types are deeply connected to immune system. HLA genes are genes in human leukocyte antigen system that is responsible for the regulation of the immune system. Genes involved in immune functions are expected to be inherently heterogeneous [26]. For example, HLA genes are highly polymorphic than others, and TR-C genes go through VDJ recombinations that lead to more diverse sequences across cells. This result corroborates with the notion that cellular heterogeneity is the main driver of zero-inflation. The high heterogeneity of cells in certain genes suggests that it is difficult, to say the least, to fully account for the cell type for every single gene; as we finely cluster the cells, we will reach a point where there are less than enough number of cells in each cluster to do statistically meaningful analysis. Still, it is practically important to resolve the cell type heterogeneity even with lower granularity, in which case we can create a blacklist of genes that expected to be more heterogeneous. These genes can be withheld from the initial analysis for a less granular, practical analysis.

**Table 3:** The alternative hypothesis *H*_*A*_ : *p*_*g*_ *> e*^−*λ*^ is robust to different model hypotheses. In the first row, the right column is larger than the left column due to Jensen’s inequality. For negative binomial, the dispersion parameter *r* is constructed so that the variance is 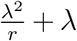, so that Poisson is a special case of Negative Binomial with *r* = *∞*. The zero-inflated negative binomial distribution is parameterized as *π*_0_*δ*_0_ + (1 − *π*_0_) NB(*λ, r*)

| Underlying Distribution | $p$ under $H_0$ | $p$ under $H_A$ |
| --- | --- | --- |
| Mixture of Poisson | $e^{-\lambda} = e^{-\sum_k \pi_k \lambda_k}$ | $\sum_k \pi_k e^{-\lambda_k}$ |
| Negative Binomial | $e^{-\lambda}$ | $\left(\frac{r}{r+\lambda}\right)^r$ |
| Zero-inflated Negative Binomial | $e^{-\lambda}$ | $\left(\frac{r}{r+\lambda}\right)^r + \pi_0$ |

**Table 4:** High gene variance is not a good indicator of cell type heterogeneity under the alternative hypothesis of zero-inflated negative binomial, because the variance can be lower under the alternative hypothesis

| Alternative Hypothesis | variance under $H_0$ | variance under $H_A$ |
| --- | --- | --- |
| Mixture of Poisson | $\lambda = \sum_k \pi_k \lambda_k$ | $\sum_k \pi_k (\lambda_k + \lambda_k^2) - (\sum_k \pi_k \lambda_k)^2$ |
| Negative Binomial | $\lambda$ | $\frac{\lambda^2}{r} + \lambda$ |
| Zero-inflated Negative Binomial | $\lambda$ | $(1 - \pi_0)^2 \left(\frac{\lambda^2}{r} + \lambda\right)$ |

### Zero inflation test for cellular heterogeneity

Based on the above observations, we propose a new feature selection strategy that uses detected zero proportion of a given gene as the statistic to test for cellular heterogeneity. Under the null hypothesis, where complete cellular homogeneity is assumed, the proportion of zeros is equal to the expected zero proportion under Poisson distribution. Under the alternative hypothesis, zero proportion is inflated, as if the count data follows mixture of Poisson (Methods). Formally, our framework can be presented as follows:

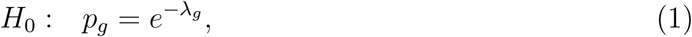

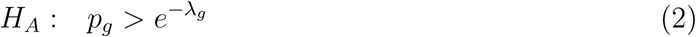

where *g* is gene index and *λ*_*g*_ is the mean UMI count for gene *g*. Above testing framework is based on an assumption that whether UMI count being 0 follows a binomial distribution. Test statistic *z*_*g*_ follows a standard normal under the null hypothesis (Methods). Genes with rejected null hypotheses will be selected for downstream analysis. For example, the CD34+ cell population within Zheng2017 dataset [36] has 2.7% of the genes with significant zero inflation at 5% Type I error level after Bonferroni correction. But after clustering into subtypes, each subtype had 1.3%, 0.3%, and 1.2% of genes with zero inflation respectively. We demonstrate the intuition of this test procedure in Table 1 using gene PPBP as an example. PPBP was identified with a high zero proportion of 26% and an average mean UMI count of 25.89 within CD34+ cells, indicating very high zero inflation with *z*-score greater than 10^6^ when the proposed test is applied. After we separate CD34+ cells into three subtypes, the test within each subtype is no longer statistically significant. We observe PPBP is highly expressed in subtype 2 and 3 and is almost unexpressed in subtype 1. This shows how cellular heterogeneity can drive excessive zeros and how zero proportions can be used to discern cell types.

Above framework significantly differs from existing ones in several ways. First of all, only the proportion of zeros (*p*_*g*_), but not that of other non-zero count values, is used in the test. We show empirically this statistic is sufficient for cellular heterogeneity analysis. This allows us not to search for a particular parametric distribution to fit all non-zero values, which can be computationally challenging. As a consequence, this feature selection strategy is robust to any data generating model and can be tied to different downstream tools (Methods). Secondly, this framework allows each gene to have different grouping structure across cells. Most existing methods select one set of genes to cluster all the cells [4, 7, 18, 34], which implicitly assume cell types can be well-defined biologically by a common set of genes. This is not realistic given the fact that each gene’s heterogeneity level varies with its function. For example, house-keeping genes are expected to behave similarly in all cells but immune-related genes, known to have more diverse genetic profiles with highly polymorphic nucleotides [14, 15], might be more finely differentiated among the sampled cells. Our approach acknowledges this type of variability. Finally, our approach provides a much more optimistic view of the UMI data analysis. No complicated modeling is needed for resolving the cellular heterogeneity.

We observe that capture efficiencies in the 10X protocols affect UMI counts in different cell types and even across datasets in a similar fashion (Supplementary Figure 1, 2). Regardless of samples or cell types, all cells show the same distribution of zero proportions with respect to mean UMI count. This means the technical noise affects each and every cell fairly, and hence, biological heterogeneity alone can largely explain the zero inflation phenomenon. Once the heterogeneity is accounted for, without any other pre-processing steps, zero proportions of UMI data closely follow the expected curve under a Poisson distribution. These observations urge us to re-evaluate some widely used pre-processing methods under this scope.

### Inappropriate pre-processing introduces unwanted noise in the downstream analysis

The most popular method for normalization is to divide UMI counts by a cell-specific scaling factor so that total UMI counts are equal across cells [32]. This strategy implicitly assumes sequencing depth effects are purely technical. However, we show sequencing depths are confounded with cell types and size factor-based adjustment can obscure biological information (Fig 2 A). For a given gene, dividing UMI counts by cell-specific factors does not change its zero proportion across cells but changes its mean. As a consequence, zero proportions across genes no longer follow the expected curve under a Poisson distribution (Fig 2) A), and the two curves from each cell type are separated from one another. For example, in 6 PBMC data sets (Azizi and Zheng) [2, 36], monocytes have lower UMI counts than B cells. The median UMI counts for monocytes and B cells, respectively, are 787 and 1180 for Zheng data for 68,000 PBMC cells, 4831 and 5575 for Azizi 2018 data for breast cancer tumors patient 9 (replication 1), 4891 and 5372 for patient 10, 5093 and 5722 for patient 11 (replication 1). When they are forced to match the median UMI count of all cells, the counts for the monocytes are inflated while those for the B cells deflated. In addition, cell types are stratified on the zero proportion plot after adjustment, indicating that total UMI counts of each cell contain valuable information about its cell type (Figure 2 A, Supplementary Figure 5). This suggests sequencing depth effects in UMI data should be accounted for differently from data generated from protocols involving PCR amplification, such as bulk RNA-seq or SMART-seq.

Sctransform is one recent influential UMI analysis method [11]. The key idea of sctransform pre-processing is to remove sequencing depth effects by introducing log-scale sequencing depth as a covariate and regressing it out from each cell under a Negative Binomial model. Similarly, this approach destroys the natural Poisson structure for zero proportions. We show in Figure 2 B an example of how normalization can further interfere with detection of biological signals. Across all 6 PBMC datasets mentioned above, we observe B cells always have more UMI counts than Monocytes before pre-processing. Applying sctransform barely modifies the sequencing depths of Monocytes but shrinks the UMI counts of B cells to match those of Monocytes. Due to the artificial shrinkage, biological markers for B cells, such as MS4A1, CD79A, and CD79B lose their power to discern B cells and Monocytes [25]. This suggests cell type differences could be potentially compromised due to excessive cleaning from sctransfrom.

Another popular pre-processing step is to apply deep learning based de-noising tools such as Deep Count Autoencoder (DCA) and SAVER, which de-convolute the technical effects from biological effects and impute zero accounts due to drop-outs at the same time. DCA implements deep neural network with flexible parametric options for noise distributions. Similarly, we observe DCA blurs the distinction among cell types, because de-noising methods essentially regularize each cell to resemble one another. We illustrate its negative impacts on downstream analysis by comparing differential expression analysis results using two imputation strategies. We selected Naive T cells and regulatory T cells from Zhengmix8eq for the experiments, which clustering algorithms often struggle to differentiate because of their similarity. In the first experiment, we imputed Naive T cells and regulatory T cells together. In the second, we imputed Naive T cells and regulatory T cells separately. Then we performed DE analysis on imputed data sets using edgeR’s likelihood ratio test [23]. We observe much greater log fold change values between Naive T and regulatory T cells from data imputed separately than data imputed together. Overall, the signal strength of DE analysis is greatly compromised across all the genes if imputing two cell types together (Fig 2 E). Using Type I error level of 0.05, 320 genes pass the Bonferroni criterion if clustering is performed first, while only 156 does if imputation is performed first. Known markers including CD4, CTLA4, FOXP3, and IL2RA [25] lost significant amount of biological signals by showing weaker log-fold change (Figure 2 D, Supplementary Figure 6). When the cells were first clustered and then imputed, the p-values were 1e-04, 8e-04, 2e-07, and 4e-11 respectively for those genes. When the cells were imputed first through DCA, the p-values were 3e-01, 4e-02, 4e-02, and 6e-07. Hence, three of the 4 genes lost statistical significance at a very liberal p-value threshold of 0.05. This analysis suggests imputing the UMI data without resolving cell heterogeneity can lead to loss of important biological information.

### HIPPO: Heterogeneity-Induced Pre-Processing Tool

The above analyses suggest the first and foremost step in pre-processing is to account for the cellular heterogeneity. Imputation or normalization before resolving the cellular heterogeneity may lead to inevitable loss of biological signals. We implement this new perspective into a computational tool called HIPPO, where we integrate the proposed zero inflation test into a hierarchical clustering framework. Specifically, we first selected genes with strong indication for cellular heterogeneity. We use a cut-off of 2 on *z*-score for selection of genes. The selected features were then used to cluster the cells into 2 groups using PCA + K-means. Then each cluster was evaluated with their intra-variability using the mean Euclidean distance from the centers of K-mean algorithm. The group with the highest intra-variability was selected and assigned for next round of clustering. The feature selection and clustering steps are iteratively repeated until one of the two ending criteria are met: *K* round of clustering for pre-determined number of clusters *K*, or the number of zero-inflated genes is less than a certain percentage of the genes. The former one can be difficult to set in real practice without any prior knowledge and the later one offers a more natural stopping criterion. HIPPO is computationally cheap because fewer and fewer features will be left for the next round of clustering, and the Poisson based test statistic has closed-form expression (Figure 3 E, F). In Figure 3 B, we show the results from each iteration of HIPPO on Zhengmix8eq data. HIPPO successfully identifies Monocytes, Natural Killer cells, B cells, and T cells in the respective order. Then it further separates naive cytotoxic cells, memory T cells, and Naive T cells from a group of Regulatory T cells and Helper T cells. However, when forced to separate into one more group, instead of clustering the remaining T cells, it created another subgroup of natural killer cells. Meanwhile, Seurat and Sctransform fails to separate the Memory T cells, Regulatory T cells, and Helper T cells, grouping them as one cluster. The adjusted rand index for the three methods show that HIPPO performs the best throughout the different K specification (Figure 3 A, D). When the selected features’ characteristics were studied through CV, gene variance, and zero proportion, Seurat and Sctransform selected more features (2000 and 3000 respectively while HIPPO selected 950), but they are highly concentrated where gene means are near 0. This is because their feature selection focuses on coefficient of variation which becomes numerically unstable as gene mean becomes near zero. HIPPO selects fewer but more relevant genes by using the zero proportion as the selection metric (Figure 3 C). This result is repeated in a different data set from muscular heart tissue in Figure 4 A. Genes selected by both methods are those with non-zero mean UMI counts, but Seurat selects extra number of genes that have mean count very close to 0. These genes are likely to add noise instead of contributing to real biological signals detection.

**Figure 3:**
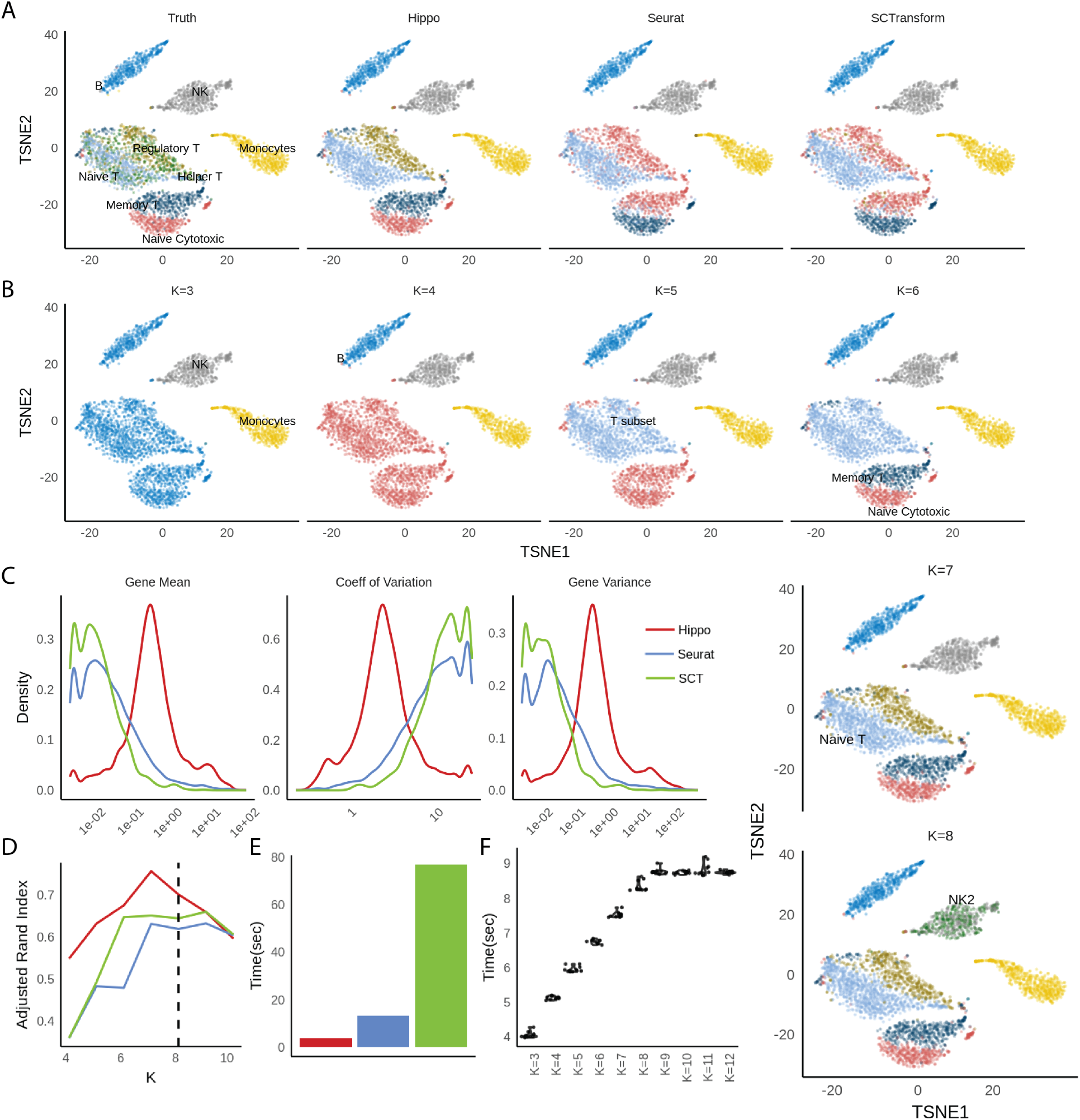
HIPPO framework applied to Zhengmix8eq data. A. t-SNE plots for clustering results from three methods: HIPPO, Seurat, and SCTransform, compared to true labels. Seurat and SCTransform cannot differentiate Helper T/Regulatory T and Memory T cells. B. HIPPO’s sequential clustering results for *K* = 3,…, 8. C. Comparisons of features selected by different methods for their gene mean, CV and variance. Seurat and SCTransform use CV as the selection criteria, and hence their features weigh heavily on genes with small mean expression and variance. D. Clustering results comparisons using Adjusted Rand Index. E. Computing time for each method using LAMBDA QUAD workstation with Intel Xeon W-2175 processor sequentially (non-parallel). F. Computing time for HIPPO using different *k*.

**Figure 4:**
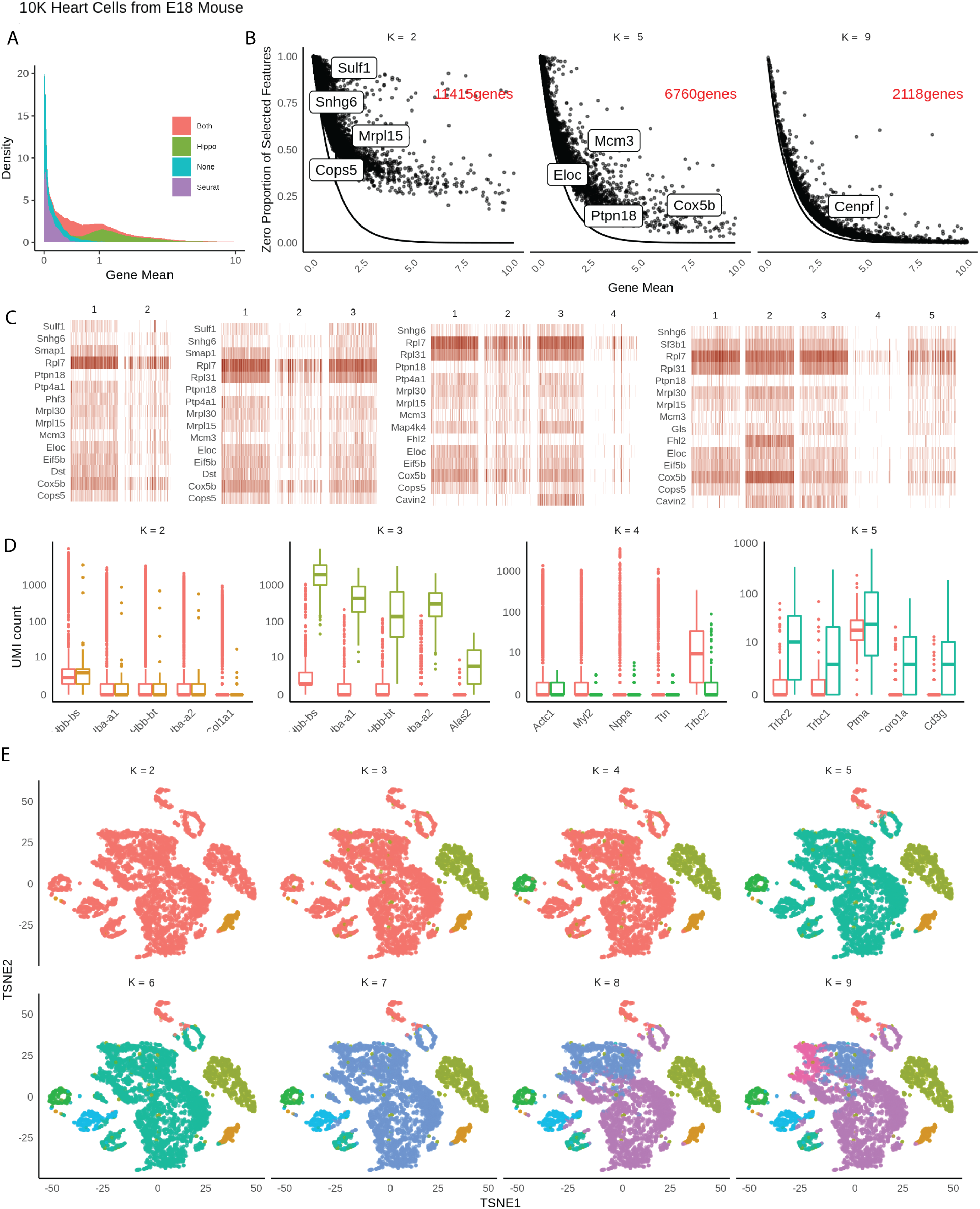
HIPPO framework applied to 10K E18 mouse heart cells. A. Distributions of means across gene features selected by Seurat, Hippo, both, or none. B. Sequential feature selection visualizes how genes gradually align closer to the expected Poisson line as more heterogeneity accounted for. C. Heatmaps of top features selected at the first 5 rounds of clustering. D. Top differential expression genes obtained at each round of clustering. E. Visualization of sequential clustering using t-SNE plots.

HIPPO’s iterative procedures naturally offer strong interpretability through sequential visualization of the analysis at each round of clustering. We use HIPPO results on an unlabeled 10X UMI data set of 10K E18 mouse heart cells for illustration. Sequential feature selection can be monitored through the visualization of the changing relationships between zero proportions and gene means. As cells are clustered into finer distinct groups, or as more cellular heterogeneity is resolved, regression lines between zero proportions and gene means get more closely aligned with the expected Poisson curve (Figure 4 B). Simultaneously, we can use a heatmap to visualize top features that contribute most at each round of clustering (Figure 4). In addition to biomarkers identified based on zero inflation, HIPPO also implements a differential expression test based on all count values to extract more features (Methods, Figure 4 D). The differential analysis can be viewed together with a t-SNE plot constructed with the same color code(Figure 4 E).

## Discussion

We have provided a new perspective on the analysis of single cell UMI data sets. Extensive analyses confirm the claims of recent literature [5] that different tool must be applied to the UMI data set from the tools for read count data set; UMI data set is free from amplification bias, so the level of technical noise is much lower. The results also show that cell-type heterogeneity must be tackled as the first step of analysis for more reliable downstream analyses. Moreover, through a streamlined feature selection method that reflects the dynamic nature of cellular process, the proposed method provides a computationally and mathematically simple analysis tool with great interpretability.

There are remaining challenges that are important in the future development of single cell UMI data analysis. First, our analysis focuses on data sets created by 10X protocols due to limited availability of labeled datasets. Still, we believe in-Drop [37] shows potential for applying the proposed modeling method (Supplementary Figure 3). In Drop-seq, the noise level was too high to assume the zero proportions follow the exponential curve relative to the gene mean (Supplementary Figure 3). It is either that Drop-seq data sets have different noise structure from the 10X data sets, or in particular Macosko data [21] of muscular retina cells have excessively high cellular heterogeneity [19]. Future new Drop-seq data could help resolve the discrepancy between 10X and Drop-seq. Secondly, although HIPPO is computationally simple compared to existing tools, the computational bottleneck is the principal component analysis, which could be slow for large cell numbers. In that case, advanced computing techniques such as sub-sampling or more rigorous filtering should be applied. Thirdly, the presented results heavily focus on PBMC samples due to data availability. In other tissues like brain, the lack of reliable cell type labels hinders our evaluation. To demonstrate wide applicability of HIPPO, we showcase analysis of muscular brain cells in Supplementary Figure 12.

We focus on the pre-processing with resolving cellular heterogeneity in our analysis tool, but this novel perspective on the noise structure of UMI data can be extended to other steps of analysis pipeline. Batch correction, lineage analysis or trajectory inference can all benefit from the simpler noise structure not only computationally but also by avoiding unnecessary normalizing steps that can introduce unwanted bias and noise.

## Methods and materials

### Datasets

Throughout the analysis, we used publicly available single cell UMI sequencing data most of which used 10X protocol. Most analysis in the main text is focused on SRP073767 which is also available in 10x Genomics, and it sequences 68,000 PBMC cells using Cell Ranger 1.1.0 [36]. We use different subsets of this data sets, namely Zhengmix4eq, Zhengmix4uneq, and Zhengmix8eq as defined in Duo (2018)[7]. Other data sets used in the main text are GSE111108 [28] and GSE115189 [9], and GSE114724 [2]. Supplementary data includes more data sets from 10X including 5k Cells from a combined cortex, hippocampus and subventricular zone of an E18 mouse (v3 chemistry), 1k Brain Cells from an E18 Mouse (v2 chemistry), and 10k Heart Cells from an E18 mouse (v3 chemistry). We also use GSE84133 [3] as an example of in-Drop and GSE63473 [21] as an example of Drop-seq. All the data sets were analyzed after their own filtering process. (2)

### Benchmarked Methods

In Figure 3, we benchmark Seurat 3.0.0 [27] and SCTransform version 0.2.0 that is integrated with Seurat platform. Seurat was implemented following its guided tutorial https://satijalab.org/seurat/v3.1/pbmc3k_tutorial.html, and SCTransform through a vignette https://rawgit.com/ChristophH/sctransform/master/inst/doc/seurat.html. All parameters were selected through software’s default except resolution parameter for clustering to generate results for various number of clusters. Seurat used in Figure 4 A was also the same version with the default parameters for feature selection. In Figure 3 A, the t-SNE plots were created using the features selected by the first round of HIPPO because they reflected the division of true cell labels the most accurately.

DCA was installed through Conda and imputation was performed following the tutorial on https://github.com/theislab/dca. In one experiment, we first divide the data set into correct labels, and then impute them separately using DCA (imputing homogeneous cell population). In the other experiment, we impute both cell types together (imputing heterogeneous cell population). One property of DCA is that it automatically removes genes that are 0 in all the cells. Naturally, there are more such genes in homogeneous cell populations. Especially, some of the biomarkers are not expressed at all when cell population is divided into subtype. In that case, we imputed zero to those genes, assuming DCA didn’t perform any imputation. (Figure 2 D, E). SAVER was downloaded from CRAN with version 1.1.1. In Supplementary Figure 7 transcriptome-level statistics were compared only using genes that had at least one positive count in each cell type. In both DCA and SAVER, all the parameters the default values as suggested by the software.

For likelihood ratio test in Figure 1 B and Figure 2 was conducted by fitting the distributions using the fitdistr function from *MASS* package [33], and zero-inflated negative binomial distribution was fitted using *pscl* package [17].

### Poisson Mixture Model

Consider a gene by cell matrix if UMI counts *X* for gene *g* = 1,…, *G* and cell *c* = 1,…, *C*. To understand the behavior of the zeros for each gene, the first step is to reduce the information from each gene to the proportion of zeros across the cells

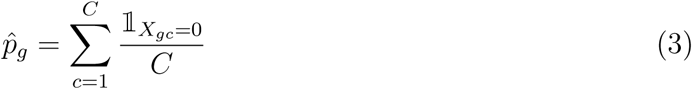

which is an estimator for the true zero proportion of gene *g*: *p*_*g*_. We study its relationship against the mean expression for the set of cells, because *p*_*g*_ would decrease as the expression level increases. With the test statistic above, we test a one-sided hypothesis for each gene *g*, whether the zero proportion is higher than the expected rate under the Poisson model. For the alternative hypothesis, we believe that UMI counts follow finite Poisson Mixture. The hypotheses for each gene *g* are formally specified below.

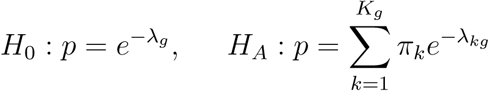

In practice, we re-frame the hypotheses as *H*_0_ : *K*_*g*_ = 1, *H*_*A*_ : *K*_*g*_ *>* 1 when 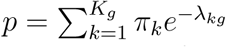. In other words, zero inflation indicates there is cell heterogeneity across the samples. If the cell population is truly homogeneous, the count data follows Poisson data with expected zero proportion 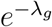.

Chen (2018) demonstrates that most genes in UMI data follow Poisson distribution [5] while other noisy genes follow Negative Binomial or Zero-Inflated Binomial distribution. Such model, although fundamentally different, is closely tied to the Poisson mixture model because Negative Binomial is the limiting distribution of Gamma-Poisson. If *λ*_*cg*_ for each cell is drawn independently from the gamma distribution 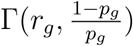, then 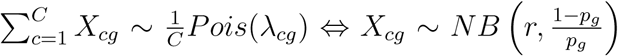. While Negative Binomial assumes a continuous mixture of Poisson, the proposed model assumes a finite mixture of Poisson, which is simpler and more directly addresses the source of zero inflation.

In practice, we do not explicitly estimate *π*_*k*_, but instead simply test if observed 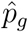 is larger than expected *p* with estimated gene mean *λ*. 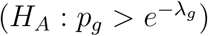. It might seem counterintuitive that this test statistic does not fully leverage the specification of the alternative hypothesis; we never estimate the mixture parameters *π*_*k*_. Alternatively, for example, one might suggest that we can conduct a likelihood ratio test of Poisson versus Poisson mixture. The main strength of the proposed reduced test statistic is its robustness to the modeling assumptions. Table 3 shows that the proportion of zeros are always larger than expected under different alternative hypotheses. Under the proposed alternative, mixture of Poisson, the proportion of zeros under the null hypothesis would be *e*^−*λ*^ where *λ* is the weighted mean of the gene mean for each cell-type. Due to Jensen’s inequality, *p* under alternative hypothesis is always greater than that under the *H*_0_.

### Feature selection and Inference

For gene *g* with count data for cells *c* = 1,…, *C*, we define an estimate for the proportion of zeros 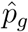. Below is our test statistic and the null hypothesis where 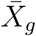 is the average counts, the maximum likelihood estimator of the true mean *λ*_*g*_.

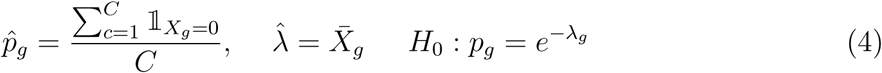

We have the following results from the above set-up.

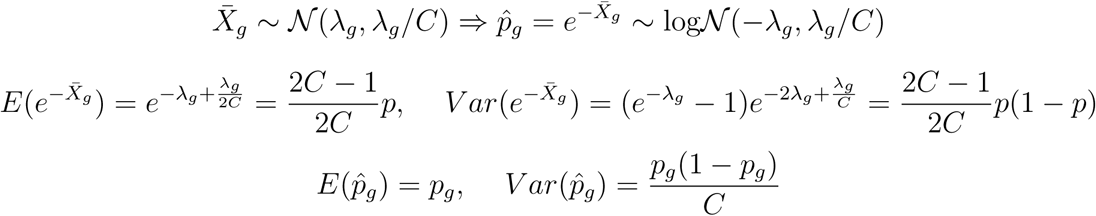

We can get the *p* value from the null distribution below

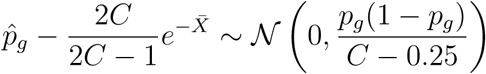

including the *z*-statistic.

### Hierarchical Clustering

Algorithm 1 outlines the iterative procedure of HIPPO’s hierarchical clustering. Several stopping criteria are determined by the user: the maximum number of clusters *K*, the feature selection statistic threshold *z*, and outlier gene proportion *o*. The algorithm first computes the number of outlier genes to allow, *G* × *o*. For example, if there are 30,000 genes in total and *o* is specified as 1% = 0.01, then the algorithm allows 300 features to have zero inflation. During the clustering procedure, HIPPO terminates in either scenarios: there are *K* identified clusters or if there are less than *G* × *o* genes that exceed the threshold. HIPPO takes all the cells and select the features whose zero inflation statistic *z* exceeds the threshold. Then it reduces the dimension through principal component decomposition using the selected features. Using the dimension-reduced cell embeddings, HIPPO clusters the cells using *K*-means. Then it selects one of the identified clusters that have the highest intra-cluster distance, and then uses that cluster for the next round of feature selection and clustering. It repeats the procedure so that each cluster in the end have the least intra-cluster distance.

#### Algorithm 1 Cell-Type Hierarchical Clustering

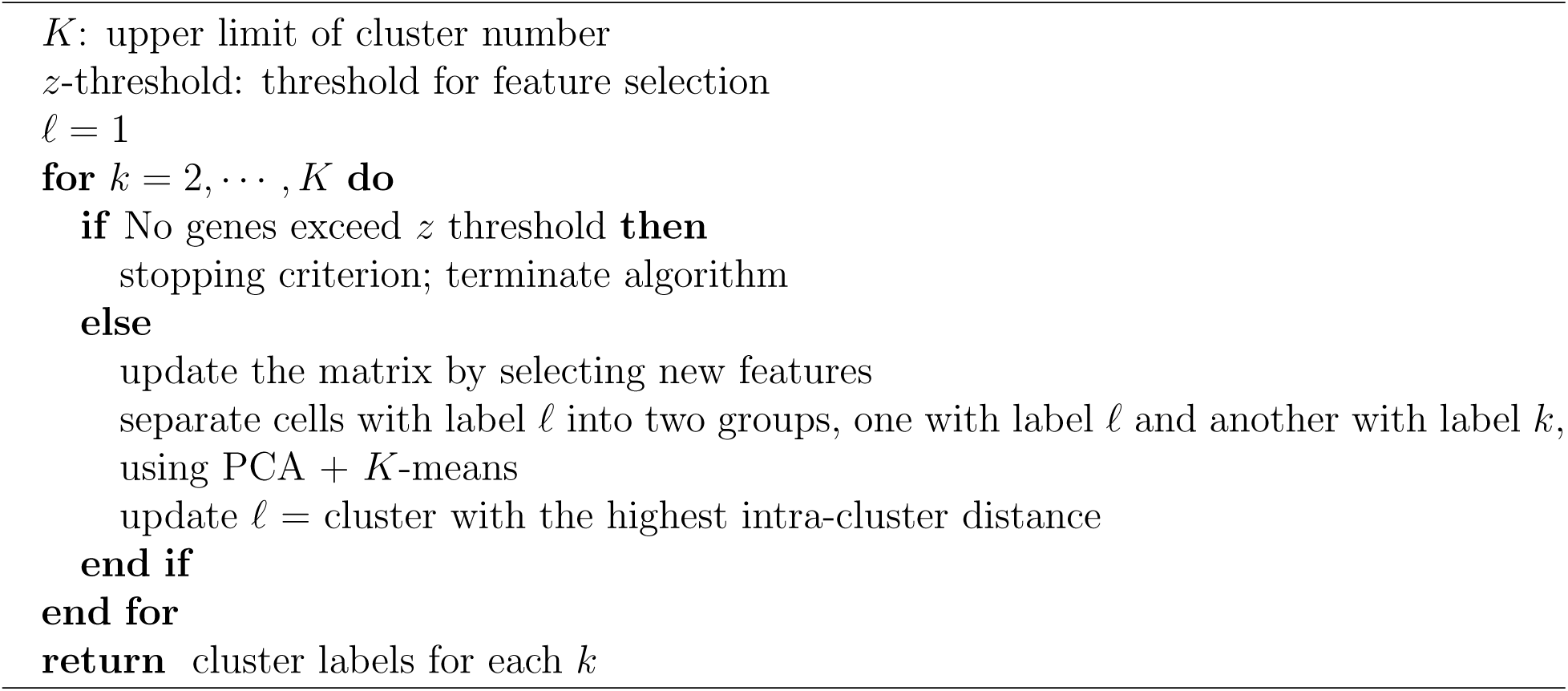

### Differential Expression Testing

After the group labeling, we can do a similar but simpler hypothesis test to see if a certain gene is differentially expressed in two groups.

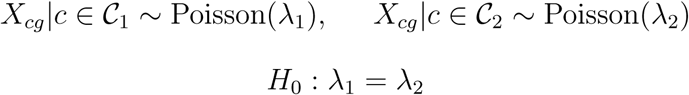

We can use a 2-sample *t*-test and order the genes in the order of significance in the mean difference of two groups.

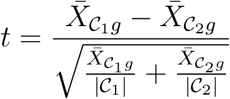

## Acknowledgment

M.C. was supported by the National Institutes of Health (NIH) Grants R01 GM126553, a Sloan Foundation Research Fellowship and a Human Cell Atlas Seed Network grant from Chan Zuckerberg Initiative. X.Z. was supported by the NIH Grants R01HG009124 and R01GM126553, and the National Science Foundation (NSF) Grant DMS1712933.

## Contributions

M.C. conceived and led this work. T.K. and M.C. developed the methods and performed the analyses. T.K. implemented the HIPPO software. X.Z. participated in critically revising the draft. T.K. and M.C. wrote the paper with feedback from X.Z.

## Competing Interests

The authors declare no competing interests.

