## Supplementary Figures and Tables for "Demystifying “drop-outs” in single cell UMI data"

**Supplementary Materials of "De-mystifying drop-outs in single cell UMI data"**

This file includes figures and tables that supplement analyses in the main manuscript.

In Section 1, we study the relationship between zero proportions and gene means in the publicly available, labeled UMI data sets of Zheng2017 and Azizi2016 (Supplementary Figures 1, 2). We explore common cell types across different datasets to emphasize that the sampling noise affects different data sets and different cell types in the same way. We study the zero proportion and gene mean relationships in Supplementary Figure 3 for data generated from other scRNAseq protocols, in-Drop and Drop-seq, to evaluate the potential of the proposed methods when applied to non-10X protocols such as Macosko2015 and Baron2016. Baron2016 (in-Drop) shows potential for applying the proposed modeling method, but Macosko2015 (Drop-seq) less so. In Drop-seq, the noise level was too high to assume the zero proportions follow the exponential curve relative to the gene mean. It is either that Drop-seq data sets have different noise structure from the 10X data sets, or in particular Macosko data of muscular retina cells have excessively high cellular heterogeneity.

In Section 2, we show why zero proportion is a better test statistic compared to gene variance and dispersion for cell-type heterogeneity using 4 data sets of Freytag2018, Zhengmix8eq, Tian2018, and Azizi2018. Supplementary Figure **??** shows the results from likelihood ratio test (Negative Binomial vs Poisson and Zero Inflated Negative Binomial vs Poisson), suggesting that the vast majority of genes can be modeled using Poisson. Supplementary Figure 4 and Supplementary Table 2 show that gene variance of homogeneous cell population is not necessarily lower than gene variance of heterogeneous cell population.

Section 3 provides the details of the analysis that show immune-related genes are more zero-inflated than others. Supplementary Table 3 shows the number of genes present within each functional annotation for each data set. Supplementary Table 4 shows the result of enrichment analysis for AziziPatient09Rep01 data set to demonstrate that zero-inflated genes are particularly enriched in immune-related genes.

In Section 4, we show additional analyses that pre-processing steps before imputation and normalization lead to adversarial consequences in downstream analyses. Supplementary Figure 5 shows that sequencing depths are confounded with the cell types, and normalization through size factors can either deflate or inflate the biological signals. Supplementary Figure 6 expands the result of Figure 2 E by showing the differential expression analysis for known markers after DCA in two cases: on homogeneous cell population and on heterogeneous population. Figure 7 shows the similar result for both DCA and SAVER but transcriptome-wide statistics for log fold change, likelihood ratio, and p-values.

Section 5 evaluates the clustering performance for more data sets: Tian2018, Zhengmix4eq, Zhengmix4uneq, Zhengmix8eq, PBMC3k1, and PBMC4k1. Supplementary Figure 8 shows the adjusted rand index for the available labeled data sets. Supplementary Figure 9 shows the sequential visualization of HIPPO's clustering method for all of those data sets. Supplementary Figure 10 evaluates the performance of generalized PCA (gPCA) that can account for the count structure directly [6].

Lastly, Section 6 includes applications of HIPPO to two different data sets of cells from muscular brain tissue (1k Brain Cells from an E18 Mouse and 5k Cells from a combined cortex, hippocampus and subventricular zone of an E18 mouse). Supplementary Figure 12 shows the clustering performance of HIPPO in two different data of un-labeled cells from brain tissue which are known to have a high level of heterogeneity. Supplementary Figure 13 shows an example analysis pipeline implemented in HIPPO. Supplementary Figure 14 visualizes the hierarchical structure of the clustering result of HIPPO through an external tool “clustree”. [10].

### 1 Supplementary Table 1: Data Sets

| ID | Data Set | Species | Protocol | Year | # Genes | # Cells |
|---|---|---|---|---|---|---|
| 10x | 5KNeuron | Mouse | 10xGenomics | 2019 | 31053 | 6997 |
| 10x | 10KHeart | Mouse | 10xGenomics | 2018 | 31053 | 7713 |
| GSE111108 [9] | Tian2018 | Human | 10xGenomics | 2018 | 58302 | 925 |
| GSE115189[4] | Freytag2018 | Human | 10xGenomics | 2018 | 58302 | 2590 |
| 10x | 1KNeuron | Mouse | 10xGenomics | 2017 | 31053 | 1206 |
| SRP073767[11] | Zhengmix4eq | Human | 10xGenomics | 2017 | 15568 | 3994 |
| SRP073767 | Zhengmix4uneq | Human | 10xGenomics | 2017 | 16443 | 6498 |
| SRP073767 | Zhengmix8efq | Human | 10xGenomics | 2017 | 15176 | 3994 |
| SRP073767 | PBMC3k | Human | 10xGenomics | 2017 | 58302 | 3250 |
| SRP073767 | PBMC4k | Human | 10xGenomics | 2017 | 58302 | 4292 |
| SRP073767 | PBMC68k | Human | 10xGenomics | 2017 | 32738 | 68579 |
| GSE84133[2] | Baron2016 | Human | inDrop | 2016 | 19097 | 350 |
| GSE114724[1] | AziziPatient09Rep1 | Human | 10xGenomics | 2016 | 33694 | 5287 |
| GSE114724 | AziziPatient09Rep2 | Human | 10xGenomics | 2016 | 33694 | 5256 |
| GSE114724 | AziziPatient10Rep1 | Human | 10xGenomics | 2016 | 33694 | 3349 |
| GSE114724 | AziziPatient11Rep1 | Human | 10xGenomics | 2016 | 33264 | 5287 |
| GSE114724 | AziziPatient11Rep2 | Human | 10xGenomics | 2016 | 33694 | 3696 |
| GSE63473[7] | Macosko2015 | Mouse | Drop-seq | 2015 | 23288 | 44808 |

Supplementary Table 1: List of data sets used in the main text and supplementary materials.

### 2 Zero Proportions in a homogeneous cell population follow a Poisson distribution.

#### 2.1 Supplementary Figure 1: Azizi 2016 across different samples

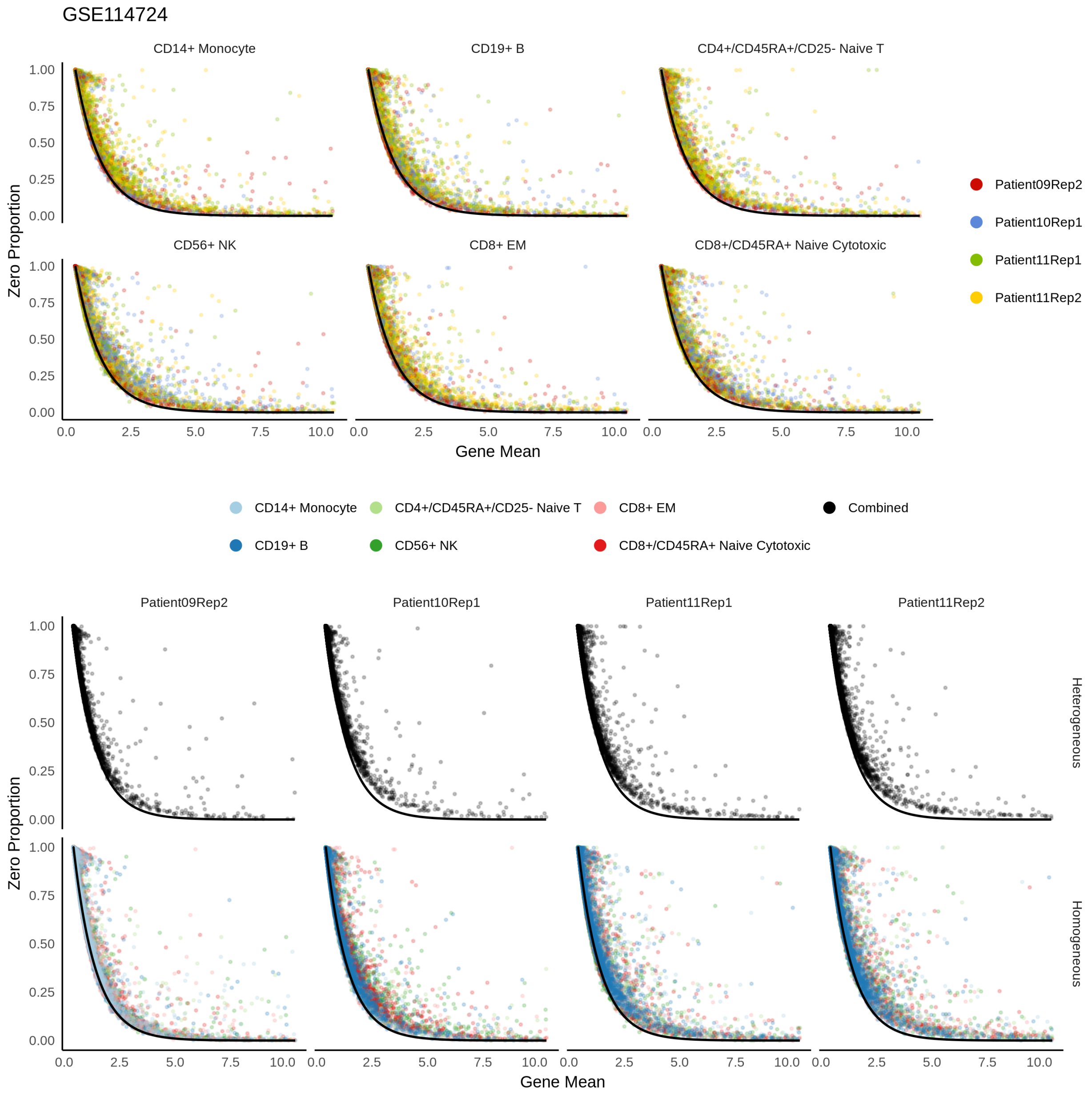

Supplementary Figure 1: Zero proportions against gene means for Azizi data [1] for multiple samples and replicates. The top plot shows that the zero proportion matches the curve across the data sets for each cell type, while bottom plot across the cell types for each data set. The bottom plot also shows that the zero proportions are off the curve in heterogeneous cell populations. The consistent plots show that the sampling noise is the same across cell types and across data sets.

#### 2.2 Supplementary Figure 2: Zheng2017 across different subsets

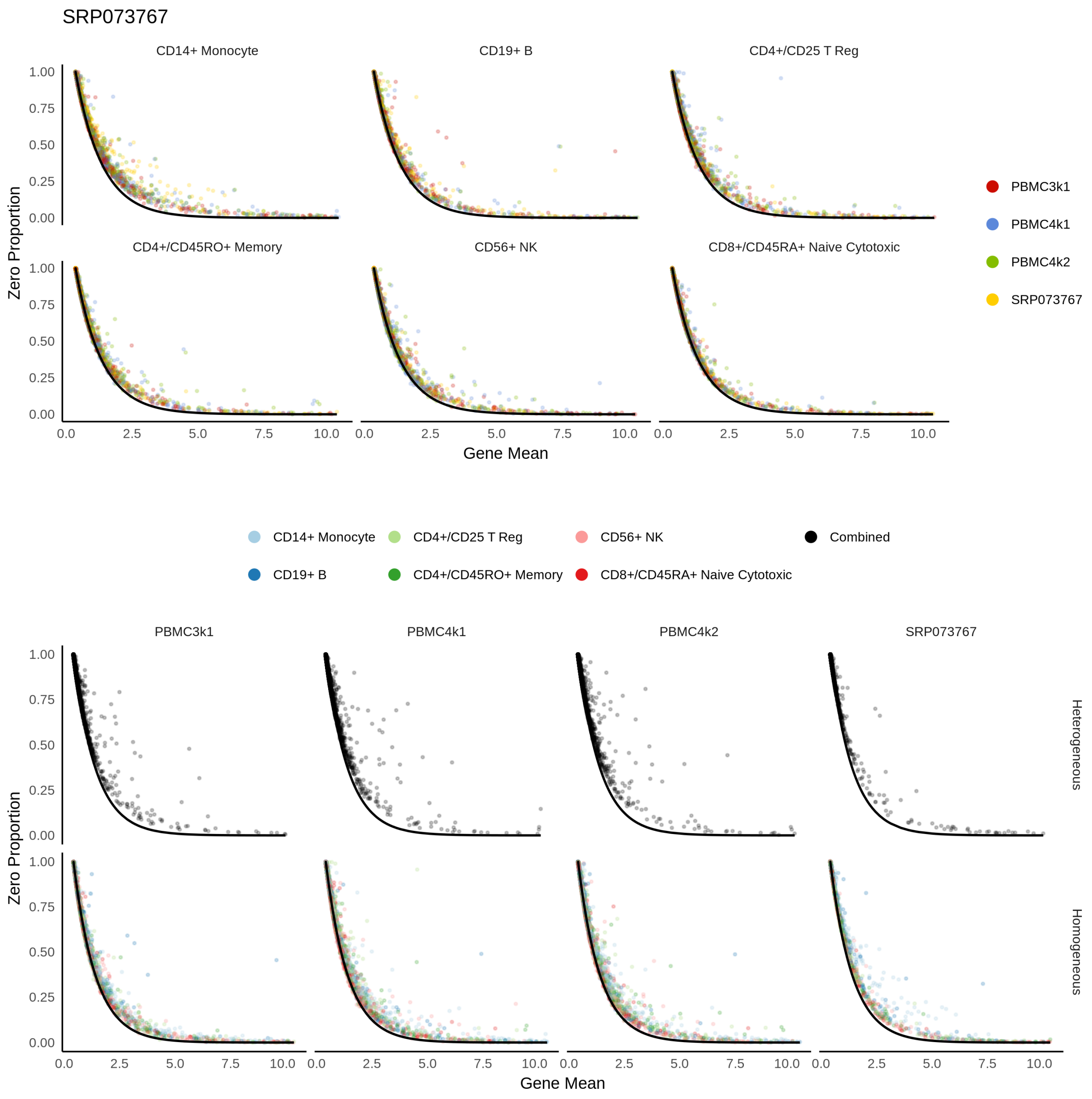

Supplementary Figure 2: Same analysis as Supplementary Figure 1 with Zheng2017 data [11]

#### 2.3 Supplementary Figure 3: Drop-Seq and In-Drop

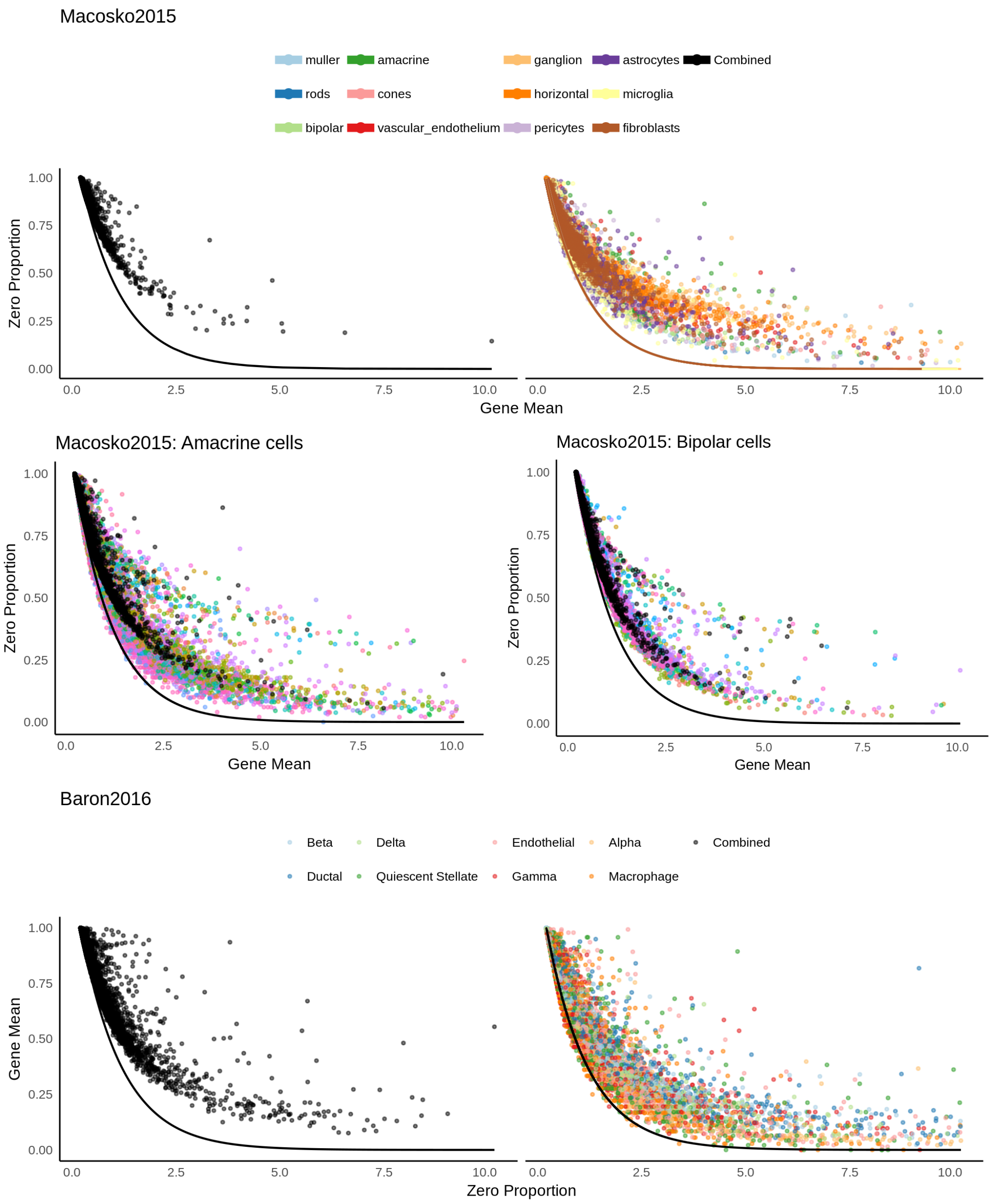

Supplementary Figure 3: Results from Drop-seq[7] and In-drop[2]. In-drop is promising that it can be modeled using Poisson. However, the sampling noise of drop-seq data is too high, and zero-inflation element seems necessary. When amacrine cells were taken out and further clustered into subtypes, the noise level is closer to Poisson, so the culprit could be the particularly higher level of cell type diversity. The black points are plotted using heterogeneous cell population.

### 3 Comparisons of zero proportions with gene variance as feature selection criteria.

#### 3.1 Supplementary Figure 4: Gene variance for heterogeneous and homogeneous Cells

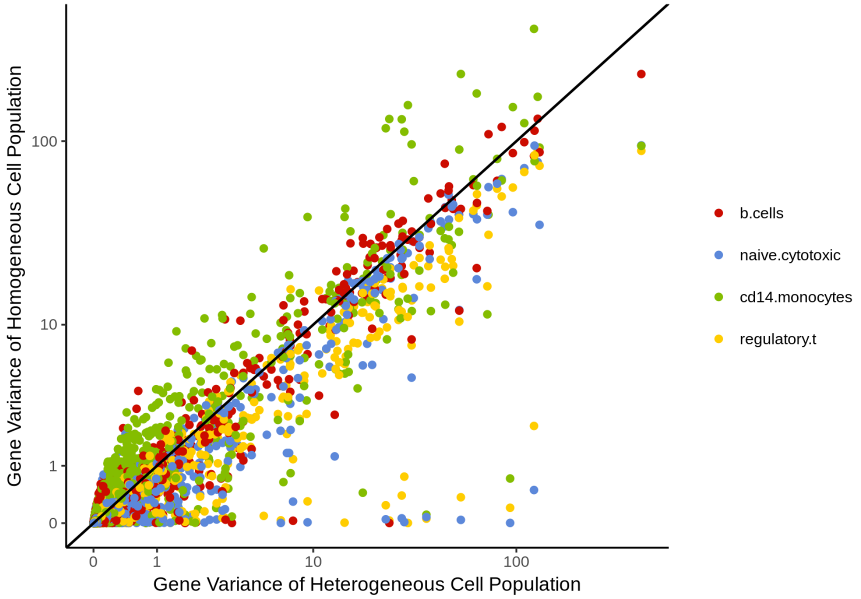

Supplementary Figure 4: Gene variance for homogeneous cell population (y axis) and heterogeneous cell population (x-axis). For most genes, gene variance is similar for both heterogeneous and homogeneous cells. Further quantifications are provided in Supplementary Table 2.

#### 3.2 Supplementary Table 2: Comparison of gene variance

| Cell Type | Proportion of Genes with Higher Variance |
|---|---|
| B cells | 0.4939857 |
| Naive Cytotoxic | 0.3700687 |
| Monocytes | 0.4969385 |
| Regulatory T | 0.4766347 |
| Helper T | 0.4620756 |
| NK | 0.7163943 |
| Memory T | 0.5348796 |
| Naive T | 0.3535812 |

Supplementary Table 2: Proportion of genes that have higher variance in heterogeneous population than in homogeneous population. Using gene variance as feature selection would not be effective for detecting cellular heterogeneity.

### 4 Immune-related genes are more zero-inflated than other functional groups.

#### 4.1 Supplementary Table 3: Blacklist genes

| | 68K | Azizi09 | Azizi10 | Azizi11 | Freytag | PBMC3k1 | PBMC4k2 |
|---|---|---|---|---|---|---|---|
| Antisense | 2356 | 1701 | 1571 | 1537 | 269 | 305 | 329 |
| HLA | 23 | 21 | 21 | 21 | 29 | 31 | 21 |
| IG-C | 0 | 9 | 11 | 10 | 13 | 13 | 13 |
| IG-C pseudo | 0 | 1 | 2 | 2 | 5 | 4 | 6 |
| IG-J gene | 0 | 0 | 1 | 3 | 1 | 2 | 0 |
| IG-V gene | 0 | 60 | 49 | 77 | 61 | 77 | 67 |
| IG-V pseudo | 0 | 6 | 6 | 5 | 1 | 7 | 6 |
| lincRNA | 2202 | 1258 | 1140 | 1170 | 183 | 210 | 216 |
| miRNA | 0 | 1 | 2 | 1 | 1 | 2 | 0 |
| misc RNA | 1 | 0 | 0 | 0 | 96 | 190 | 0 |
| Mt-rRNA | 0 | 0 | 0 | 0 | 2 | 2 | 0 |
| MT-tRNA | 0 | 0 | 0 | 0 | 12 | 11 | 0 |
| Polymorphic pseudo | 0 | 6 | 6 | 6 | 11 | 11 | 7 |
| Processed transcripts | 3 | 58 | 52 | 55 | 86 | 88 | 46 |
| Protein coding | 15338 | 13684 | 13493 | 14080 | 12326 | 13025 | 13347 |
| rRNA | 0 | 0 | 0 | 0 | 14 | 18 | 0 |
| Sense intronic | 0 | 4 | 5 | 4 | 31 | 51 | 1 |
| Sense overlapping | 0 | 1 | 1 | 1 | 3 | 3 | 0 |
| snoRNA | 2 | 0 | 0 | 0 | 17 | 38 | 0 |
| snRNA | 0 | 0 | 0 | 0 | 71 | 134 | 0 |
| TR-C gene | 0 | 5 | 5 | 5 | 5 | 5 | 5 |
| TR-J gene | 0 | 1 | 2 | 46 | 0 | 0 | 0 |
| TR-V gene | 0 | 92 | 90 | 90 | 69 | 76 | 80 |
| TR-V pseudogene | 0 | 12 | 10 | 9 | 4 | 3 | 5 |

Supplementary Table 3: Gene counts for each data set and each gene type for PBMC data [1, 4, 11]. Most of the genes are categorized as protein coding genes.

##### 4.2 Supplementary Table 4: Enrichment Analysis

| Azizi Patient 9 Rep1 | | $z \leq 3$ | $z > 3$ | Proportion | $\chi_1^2$ statistic |
|---|---|---|---|---|---|
| Before Clustering | Immune genes | 98 | 109 | 52.66% | 553.66 |
| | Others | 15476 | 1262 | 7.54% | $p < 2.2e - 16$ |
| After Clustering | Immune Genes | 826 | 678 | 45.08% | 10960 |
| | Others | 145941 | 5152 | 3.41% | $p < 2.2e - 16$ |

Supplementary Table 4: Azizi Patient9 Replication 1. Immune-related genes include HLA-gene, IG C gene, IG C pseudogene, IG V gene, IG V pseudogene, TR C gene, TR J gene, TR V gene, and TR V pseudogenes. The $\chi_1^2$ statistic is computed through Pearson's chi squared test for independence of the two by two table. Clustering was performed using the true labels provided by the original paper [1]. Each gene is recorded once for each cell type, explaining the increase of the number of genes. By repeating the Pearson's chi squared test for the combined data for each cell type, we are implicitly assuming that each cell types are independent.

### 5 Unwanted consequences of pre-processing when cell-type heterogeneity is not appropriately accounted for.

#### 5.1 Supplementary Figure 5: Sequencing Depths

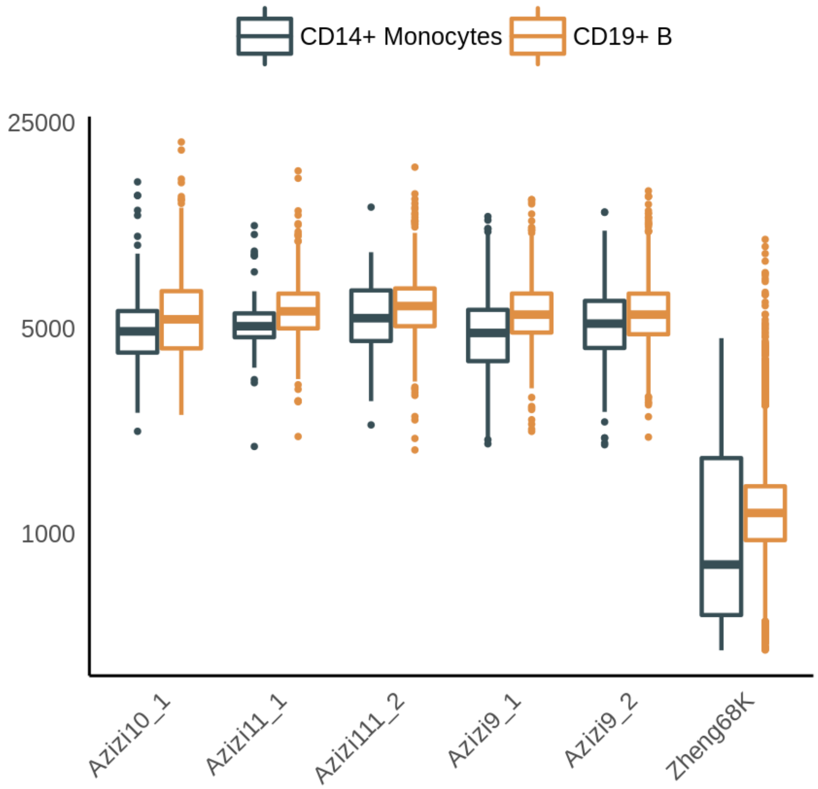

Supplementary Figure 5: Sequencing depth of monocytes and B cells. Monocytes have consistently higher total UMI counts than B cells, and forcing all the cells to have the same sequencing depth (size factor normalization) would either shrink the counts of B cells or inflate the counts of monocytes.

#### 5.2 Supplementary Figure 6: Imputation

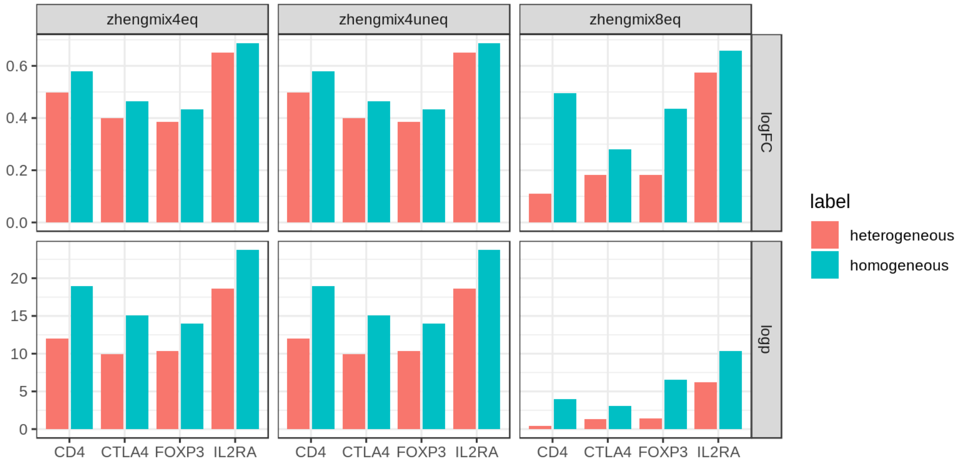

Supplementary Figure 6: Extension of Figure 2 E in the main text. The log-fold change is consistently lower across data sets if DCA [3] is performed before the cell heterogeneity is accounted for.

#### 5.3 Supplementary Figure 7: Distribution of statistics

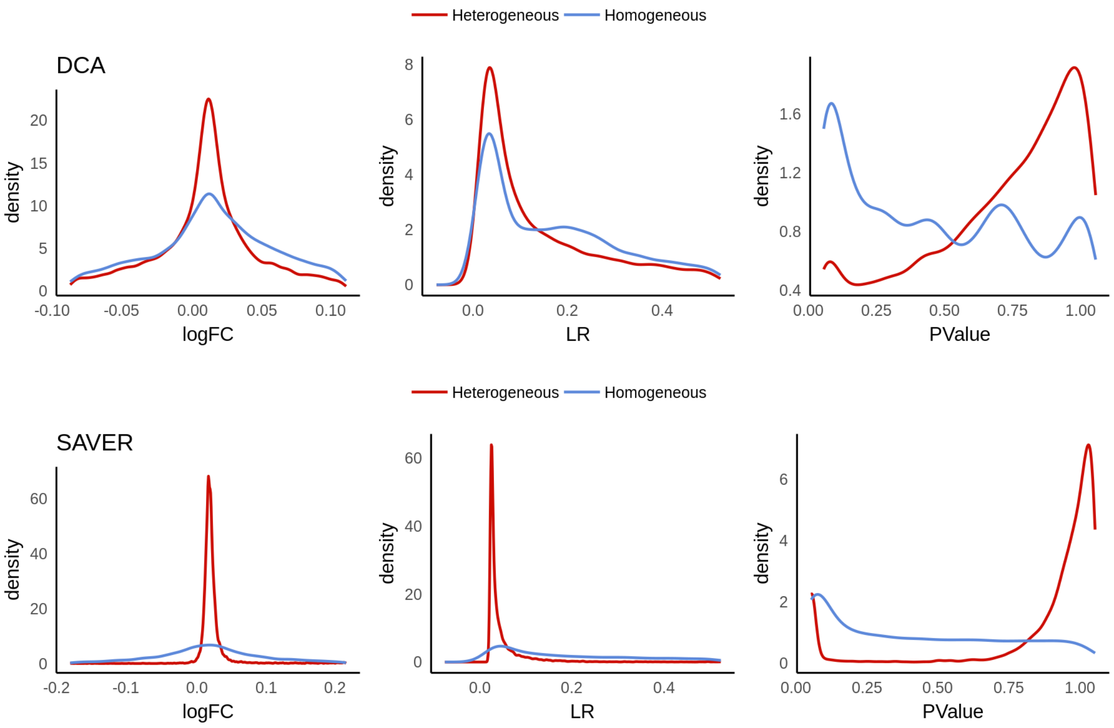

Supplementary Figure 7: Extension of Supplementary Figure 6. Overall distribution of various statistics (log fold change, likelihood ratio, and p-value) from differential expression test using edgeR's likelihood ratio test [8] after DCA [3] and SAVER [5]. Overall signal size is deflated if we perform imputation first.

### 6 Comparisons of clustering performance using different tools

#### 6.1 Supplementary Figure 8: ARI comparison with Seurat and Sctransform

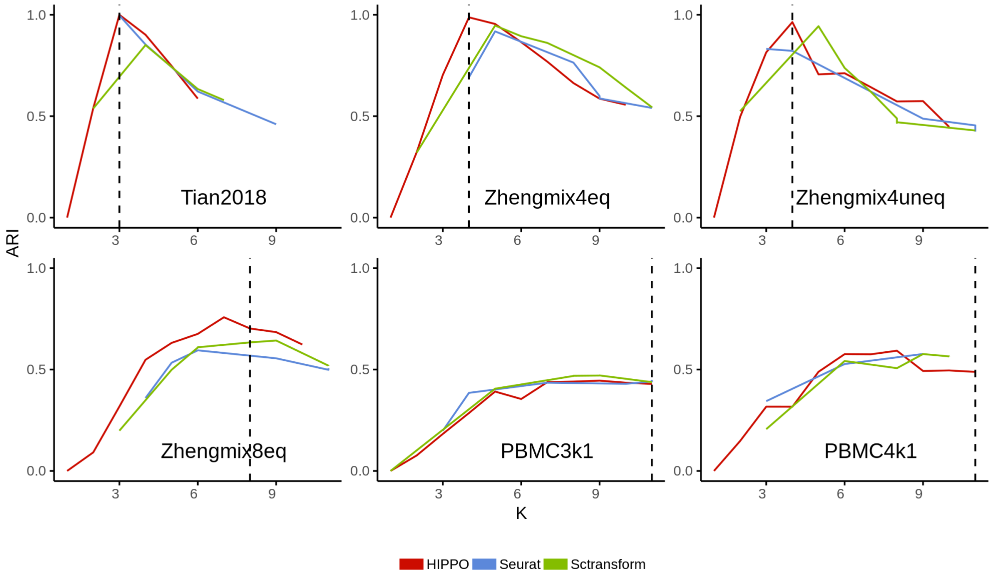

Supplementary Figure 8: Adjusted Rand Index for various data sets comparing three methods. HIPPO tends to work at least as well as Sctransform and Seurat.

#### 6.2 Supplementary Figure 9: Sequential visualization

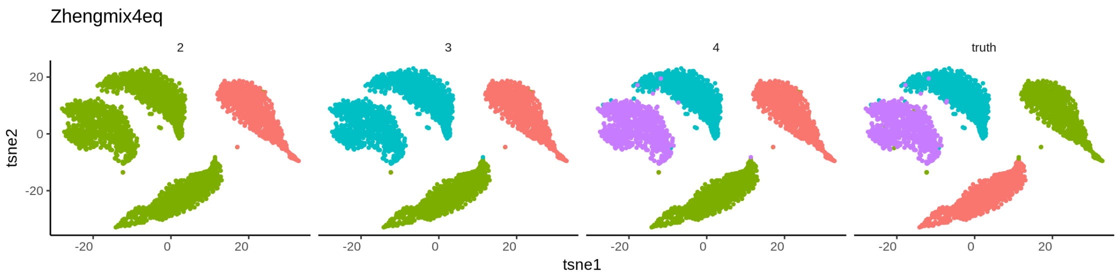

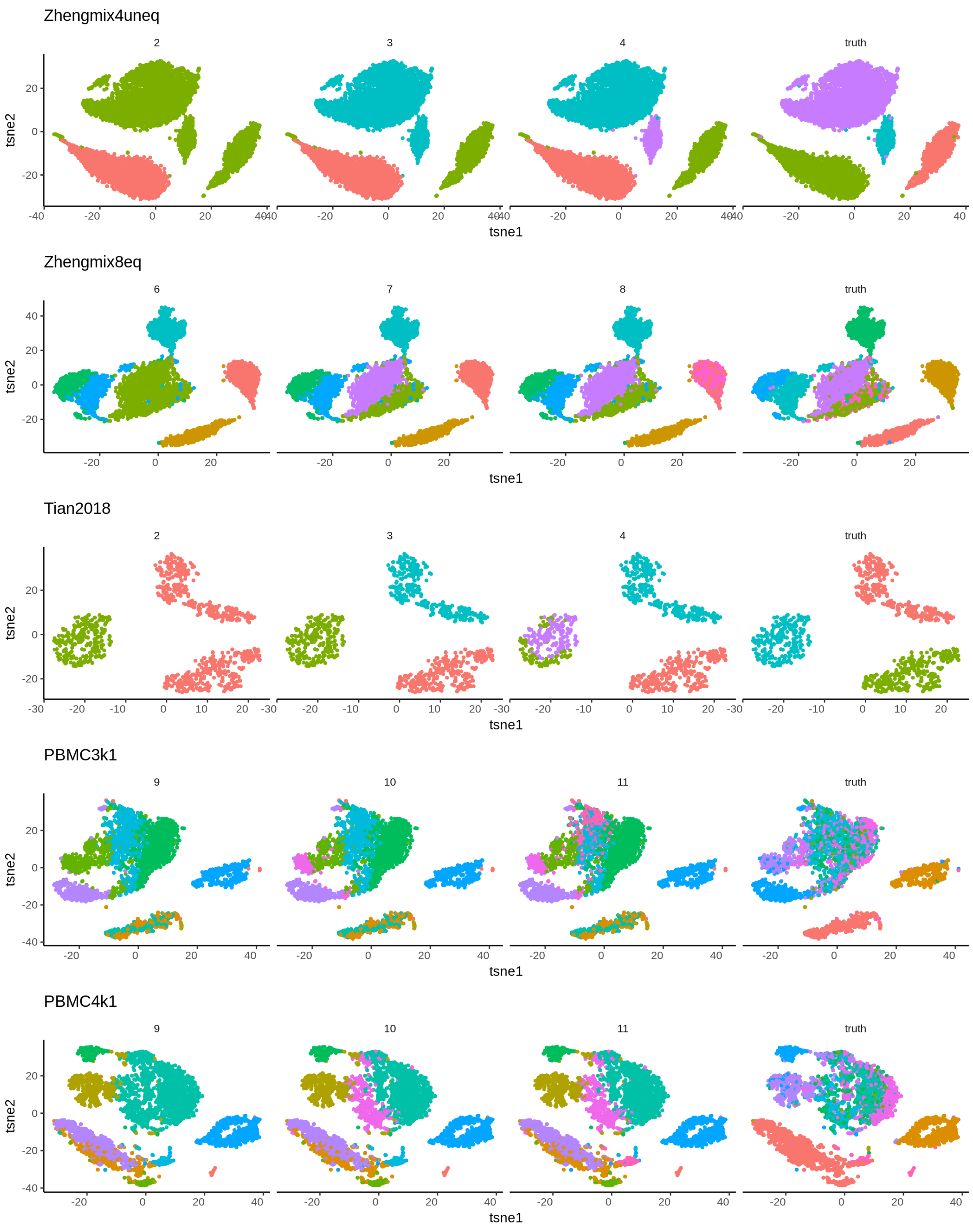

Supplementary Figure 9: Visualization of the step-by-step clustering of HIPPO in various data sets. One drawback is that when it can no longer identify distinct clusters and forced to cluster into more groups, it can divide existing groups into subsets and drive down the adjusted rand index.

#### 6.3 Supplementary Figure 10: Generalized PCA

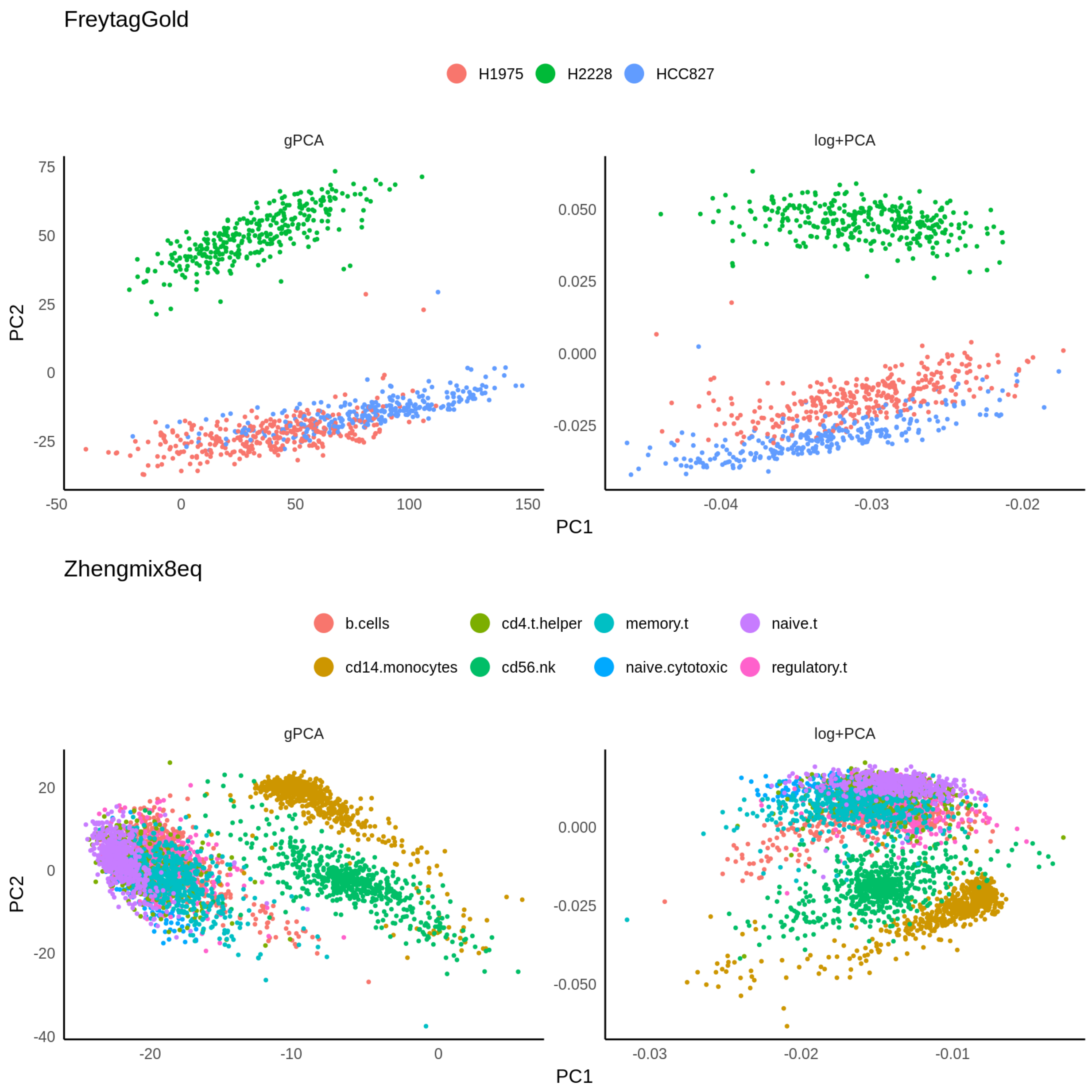

Supplementary Figure 10: Generalized PCA (gPCA) [6] takes into acount the count structure of the data to reduce the dimensions, and could be integrated into HIPPO procedure. However, empirically, its results are similar to the result of log transformation + PCA, and the result does not make up for the computational burden of gPCA.

### 7 Analysis with HIPPO

#### 7.1 Supplementary Figure 11: HIPPO applied to Brain cells

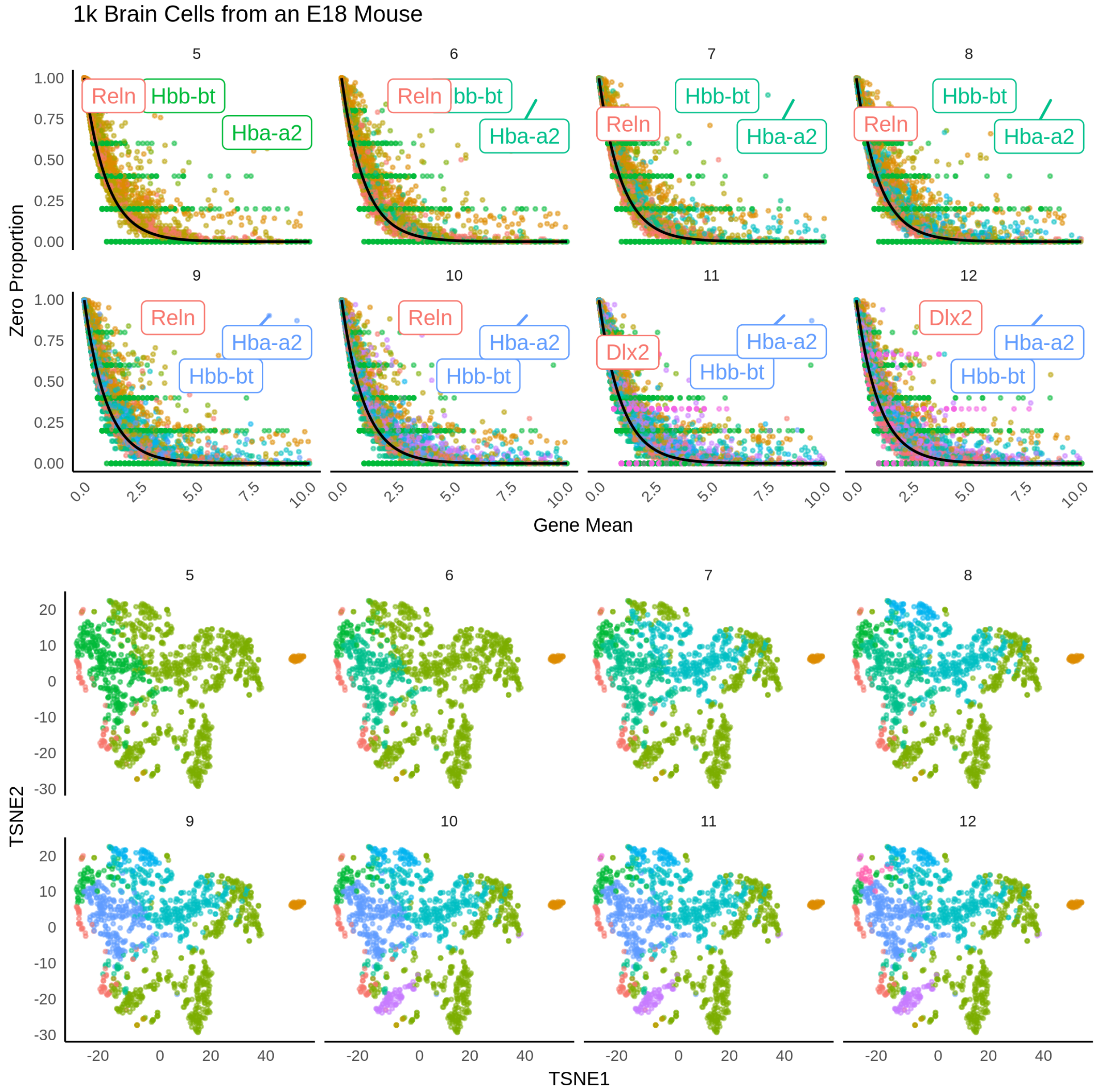

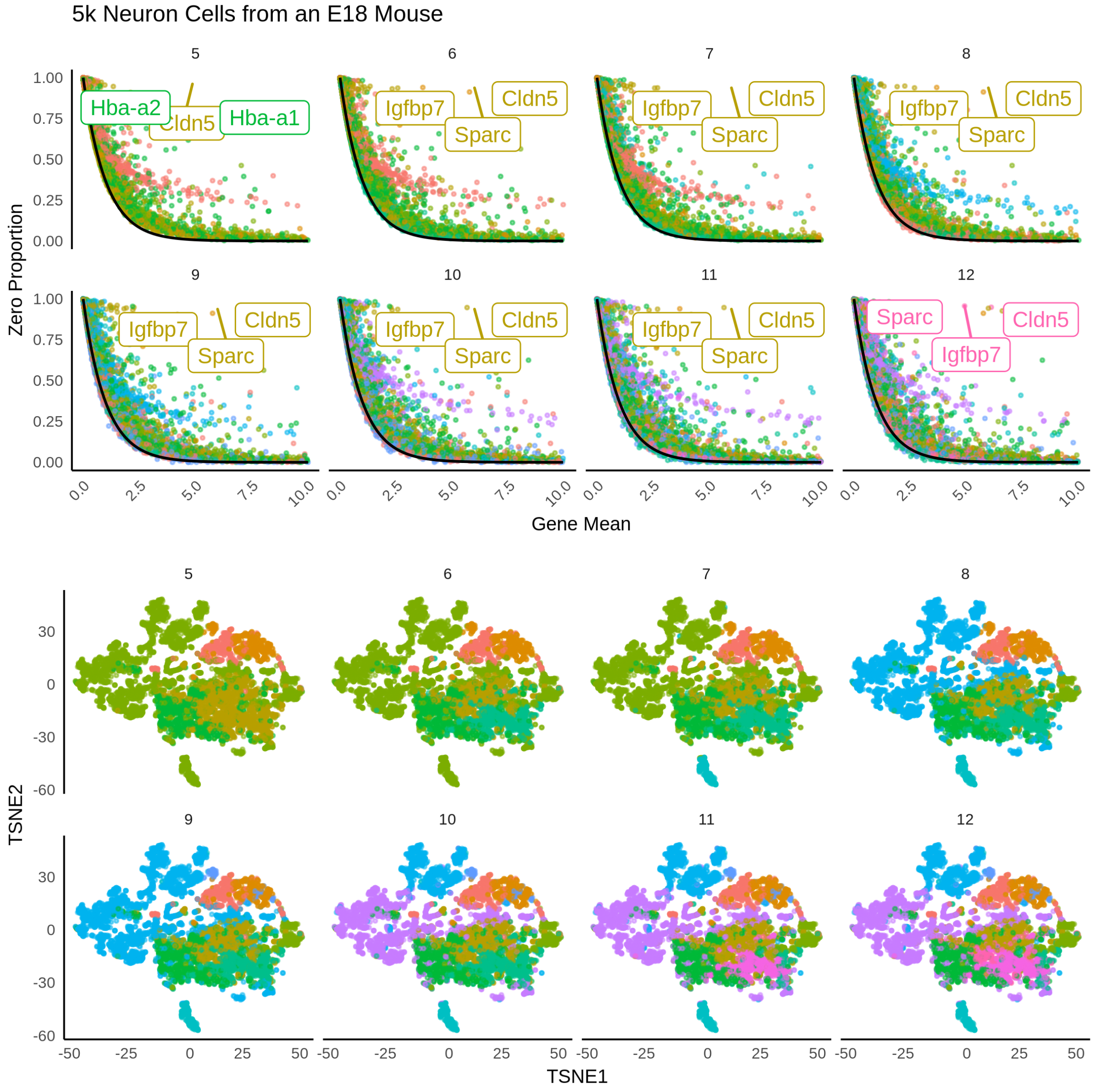

Supplementary Figure 11

Supplementary Figure 12: Example analysis of HIPPO for cells from two examples of brain tissues with higher number of clusters. For each round of clustering, zero proportions are more aligned to the Poisson line. The t-SNE plot is more finely separated as the number of clusters increase. HIPPO can show the differentiation of cell types in sequencing manner.

#### 7.2 Supplementary Figure 12: HIPPO Analysis pipeline

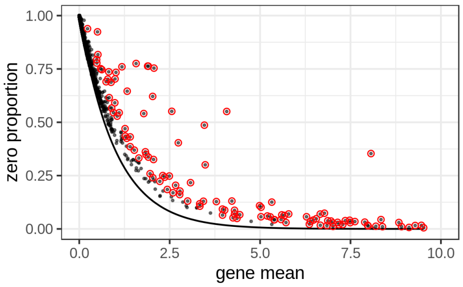

(a)

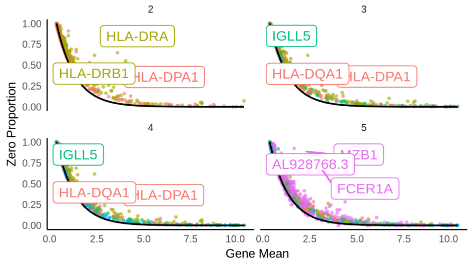

(b)

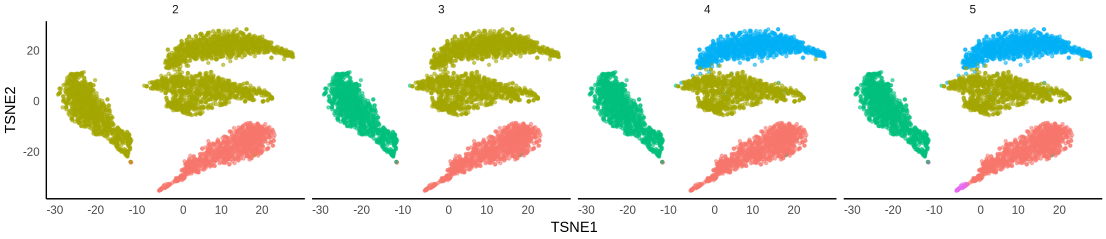

(c)

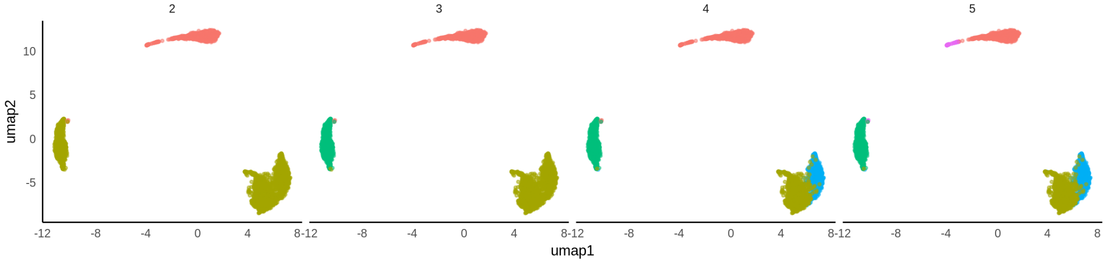

(d)

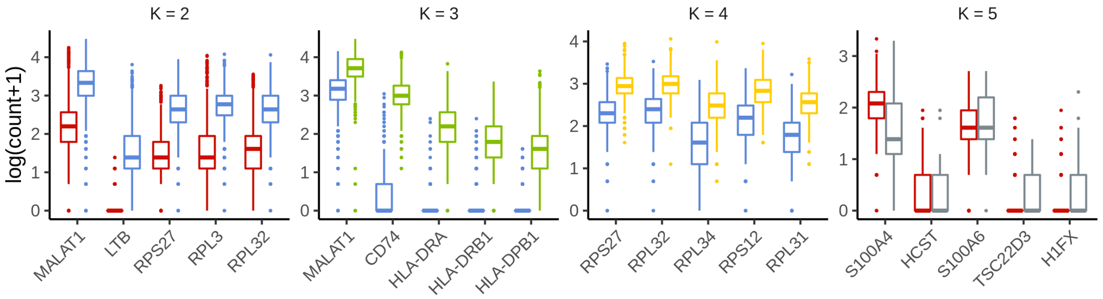

(e)

Supplementary Figure 13: Sample analysis of Zhengmix4eq using HIPPO. The software first shows the diagnostic plot where zero-inflated genes are marked in red. Then it performs the clustering which leads to three sequential plots: zero proportions, t-SNE, and UMAP. Lastly, it shows the sequential differential expression analysis where color-coding matches the t-SNE and UMAP plots.

#### 7.3 Supplementary Figure 13: Tree structure of Hierarchical Clustering

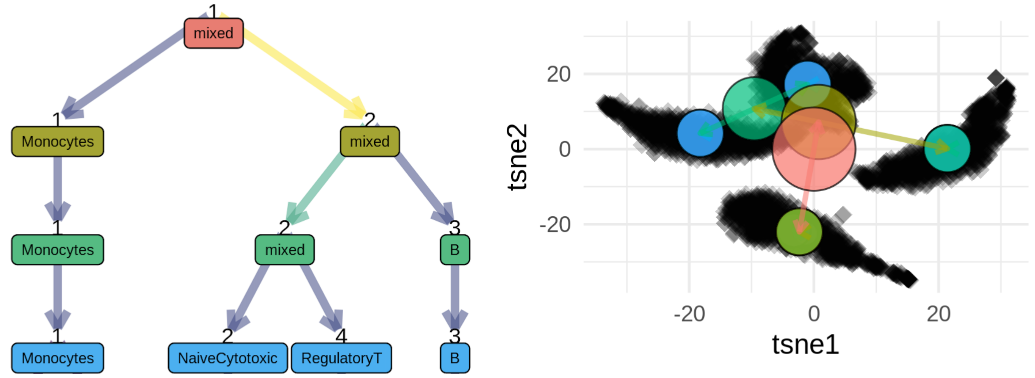

Supplementary Figure 14: Clustree [10] package allows the tree-like visualization of the clustering result. The hierarchical structure gives insight to the overall structure of cell types and subtypes.

[4] Saskia Freytag, Luyi Tian, Ingrid Lönnstedt, Milica Ng, and Melanie Bahlo. Comparison of clustering tools in r for medium-sized 10x genomics single-cell rna-sequencing data. *F1000Research*, 7, 2018.

[5] Mo Huang, Jingshu Wang, Eduardo Torre, Hannah Dueck, Sydney Shaffer, Roberto Bonasio, John I Murray, Arjun Raj, Mingyao Li, and Nancy R Zhang. Saver: gene expression recovery for single-cell rna sequencing. *Nature methods*, 15(7):539, 2018.

[6] Yoonkyung Lee. Generalized principal component analysis. *Journal of Educational Psychology*, 24(6):417–441, 2015.

[7] Evan Z Macosko, Anindita Basu, Rahul Satija, James Nemesh, Karthik Shekhar, Melissa Goldman, Itay Tirosh, Allison R Bialas, Nolan Kamitaki, Emily M Martersteck, et al. Highly parallel genome-wide expression profiling of individual cells using nanoliter droplets. *Cell*, 161(5):1202–1214, 2015.

[8] Mark D Robinson, Davis J McCarthy, and Gordon K Smyth. edger: a bioconductor package for differential expression analysis of digital gene expression data. *Bioinformatics*, 26(1):139–140, 2010.

[9] Luyi Tian, Xueyi Dong, Saskia Freytag, Kim-Anh Le Cao, Shian Su, Daniela Amann-Zalcenstein, Tom S Weber, Azadeh Seidi, Shalin Naik, and Matthew E Ritchie. scrna-seq mixology: towards better benchmarking of single cell rna-seq protocols and analysis methods. *BioRxiv*, page 433102, 2018.

[10] Luke Zappia and Alicia Oshlack. Clustering trees: a visualization for evaluating clusterings at multiple resolutions. *GigaScience*, 7(7), jul 2018.

[11] Grace XY Zheng, Jessica M Terry, Phillip Belgrader, Paul Ryvkin, Zachary W Bent, Ryan Wilson, Solongo B Ziraldo, Tobias D Wheeler, Geoff P McDermott, Junjie Zhu, et al. Massively parallel digital transcriptional profiling of single cells. *Nature communications*, 8:14049, 2017.
